# Bassoon is required for adult mouse ocular dominance plasticity and experience-induced changes in synapsin phosphorylation in visual cortex

**DOI:** 10.64898/2026.09.21.753144

**Authors:** Cornelia Schöne, Carolina Montenegro-Venegas, Debarpan Guhathakurta, Bianka Barthel, Franziska Greifzu, Josephin Böhner, Santosh Pothula, Eneko Pina-Fernandes, Anna Beth, Anil Annamneedi, Martin Heine, Eckart D. Gundelfinger, Siegrid Löwel, Anna Fejtova

## Abstract

Sensory stimulation enables experience-dependent fine tuning of brain circuits for optimal perception. In juvenile mice, brief closure of the contralateral eye reduces primary visual cortex (V1) activation through the deprived eye. In contrast, in adult animals, eye closure induces potentiated V1-responses to non-deprived eye visual stimulation. While considerable knowledge exists about postsynaptic mechanisms underlying this ocular dominance (OD) plasticity, little is known about contributing presynaptic molecules. Here we report that constitutive deletion of the presynaptic scaffolding protein bassoon (Bsn^ΔEx4/5^) disrupted OD-plasticity in V1 of adult mice leaving juvenile OD-plasticity unaffected. Notably, in mice with a conditional deletion of *Bsn* only from cortical glutamatergic neurons (Bsn^Emx1^), adult OD-plasticity was also impaired indicating that normal presynaptic plasticity of glutamatergic cortical synapses is selectively required in this process. While spatial vision was severely compromised in Bsn^ΔEx4/5^ mice, it was preserved in Bsn^Emx1^ mice, consistent with normal Bsn expression in subcortical circuit in these mice. Parallel experiments in cultured neurons revealed that deletion of *Bsn* left the inactivity-induced postsynaptic scaling of AMPA-responses mediated by increased AMPA receptor trafficking unchanged. However, presynaptic scaling was completely abolished as assessed by both electrophysiology and live-cell imaging of evoked synaptic vesicle fusion in cultured primary cortical neurons. Finally, monocular deprivation induced enhanced phosphorylation of synaptic vesicle-associated synapsin in contralateral V1 that was absent in Bsn^Emx1^ mice. Overall, our data indicate that adult OD-plasticity requires *Bsn*-dependent presynaptic plasticity, involving experience-induced changes in synapsin phosphorylation. In contrast, juvenile OD-plasticity does not appear to be dependent on *Bsn*.

## Introduction

The visual cortex is a prime model system to test the physiological and perceptual consequences of modified sensory experience. In the binocular region of the mouse primary visual cortex (V1), stimulation of the contralateral eye elicits a much stronger response than stimulation of the ipsilateral eye (Dräger, 1975; Gordon and Stryker, 1996; Porciatti et al., 1999; Coleman et al., 2009). Unbalanced sensory experience, like monocular vision after brief monocular deprivation (MD), leads to a rapid shift of V1-responses in favor of the open, experienced eye (Drager, 1978; Gordon and Stryker, 1996; Sawtell et al., 2003; Cang et al., 2005; Sato and Stryker, 2008). The magnitude and velocity of this OD-plasticity primarily depend on age, housing and MD-duration (Lehmann and Löwel, 2008; Espinosa and Stryker, 2012; Ranson et al., 2012; Greifzu et al., 2014; Hosang et al., 2018; Stryker and Löwel, 2018). In juvenile mice during the critical period (CP) for OD-plasticity, 4 days of MD are sufficient to induce an OD-shift, mediated by reduced V1-activation after visual stimulation of the previously deprived eye. In contrast, in 2-3 months old standard-cage raised mice, OD-shifts require at least 7 days of MD and are mainly driven by increased V1-activation via visual stimulation of the open eye (Lehmann and Löwel, 2008; Sato and Stryker, 2008). Adult OD-plasticity is N-methyl-D-aspartic acid (NMDA) receptor-dependent and likely involves input-specific as well as homeostatic plasticity mechanisms (Sawtell et al., 2003; Hensch and Quinlan, 2018; Stryker and Löwel, 2018). Within the cortex, the neuronal loci for this plasticity are excitatory cells of the thalamic input layer 4, layer 2/3 neurons as well as inhibitory interneurons (Desai et al., 2002; Sawtell et al., 2003; Hensch, 2005; Goel and Lee, 2007; Kuhlman et al., 2013; Lambo and Turrigiano, 2013; Stryker and Löwel, 2018; Wen and Turrigiano, 2021).

At the molecular level, the input-specific and homeostatic mechanisms implicated in OD-plasticity show a high degree of overlap. They include alterations in surface expression of NMDA and alpha-amino-3-hydroxy-5-methyl-4-isoxazolepropionic acid (AMPA) receptors (AMPARs), induction of signaling by protein kinase A (PKA) mitogen-activated kinase (MAPK), calmodulin-activated kinase type II (CaMKII), and changes in gene expression (Tagawa et al., 2005; Tropea et al., 2009). While considerable knowledge exists about postsynaptic mechanisms of OD-plasticity, presynaptic mechanisms have been largely ignored, despite the fact that sensory deprivation in V1 was shown to induce changes in presynaptic strength and that major signaling cascades implicated in OD-plasticity are also active in the presynaptic compartment (Keck et al., 2013). Supporting a role of the presynapse for OD-plasticity, expression of a presynaptic constitutively active form of HRAS proto-oncogene, a signaling molecule involved in regulating synaptic plasticity, accelerated the rate of MD-induced OD-plasticity at least partially by modulating efficacy of neurotransmitter release from the presynaptic ending (Kaneko et al., 2010).

Bassoon (BSN) is expressed in virtually all neurons and localized exclusively to their presynaptic endings, where it plays a critical role in the structural and functional organization of presynaptic release sites (Gundelfinger and Fejtova, 2012; Gundelfinger et al., 2015). Deletion of the *Bsn* gene reduces the release competence of synaptic vesicles (SVs) limiting their ability to undergo evoked fusion (Montenegro-Venegas et al., 2022). This leads to an increase in presynaptically silent synapses, lower evoked neurotransmitter release and stronger depression of neurotransmission upon repetitive stimulation (Altrock et al., 2003; Hallermann et al., 2010). At the molecular level, *Bsn* controls the balance between presynaptic phosphorylation and dephosphorylation (Montenegro-Venegas et al., 2022). *Bsn* deletion reduces presynaptic activity of the cyclic adenosine monophosphate (cAMP) /PKA pathway and shifts the cyclin-dependent kinase 5 (CDK5)/calcineurin signalling resulting in an aberrant phosphorylation of the SV-associated protein synapsin (SYN), which in turn causes defective neurotransmitter release (Montenegro-Venegas et al., 2022). Interestingly, the cAMP/PKA pathway and CDK5/calcineurin balance have been implicated in MD-induced OD-plasticity, but so far mostly studied in the context of postsynaptic regulations (Tropea et al., 2009; Li et al., 2018). While changes in the phosphorylation of SYN during development or in stimulus-dependent manner have already been described in the visual cortex, underlying mechanisms have not yet been addressed (Scott et al., 2010; Fu et al., 2015)..

Previous studies using multiple mouse models - including the Bsn^ΔEx4/5^, Bsn^GT^ and Bsn^KO^ mice used here - revealed that lack of functional *Bsn* affects the morphology of the ribbon synapses of retinal photoreceptors and bipolar cells that impaired ribbon attachment to the active zone and caused a severely disturbed signal transfer at photoreceptor ribbon synapses and compromised light-induced retinal responses in these animals (Dick et al., 2003; Babai et al., 2019; Babai et al., 2021; Ryl et al., 2021). Additionally, a progressive retinopathy with a cone photoreceptor cell-death and sprouting of bipolar and horizontal cell neurites developed in adult Bsn^ΔEx4/5^ and Bsn^GT^ mice (Specht et al., 2007; Ryl et al., 2021). Bsn^ΔEx4/5^ mice showed drastically reduced visual acuity, but were still able to discriminate visual cues in a dual-choice visual discrimination task (Goetze et al., 2010). While spatial vision was severely compromised, as documented by both substantially diminished contrast sensitivity and spatial frequency thresholds (SFT) of the optomotor reflex, visually evoked V1-activity and retinotopic maps visualized by intrinsic signal optical imaging were indistinguishable from wildtype controls (Bsn^WT^) in both juvenile and adult animals in spite of the severe photoreceptor synaptopathy (Goetze et al., 2010). In the present study, we aimed to investigate the importance of *Bsn* expression for experience-dependent changes in the visual cortical networks and specifically, whether *Bsn*-dependent presynaptic plasticity involving the regulation of SYN phosphorylation can contribute to OD-plasticity in mouse V1.

Therefore, we tested OD-plasticity in juvenile and adult mice with a global deletion of functional Bassoon (Bsn^ΔEx4/5^). In addition, we tested OD-plasticity also in Bsn^WT^ littermates and measured spatial vision before and after MD, using optometry. Since constitutive *Bsn* mutant mice suffer from epileptiform seizures involving the cerebral cortex (Altrock et al., 2003; Ghiglieri et al., 2009; Ghiglieri et al., 2010; Sgobio et al., 2010; Blondiaux et al., 2023) which might interfere with OD-plasticity, we additionally analyzed a constitutive *Bsn* mutant (Bsn^Emx1^) mouse line with restricted ablation of *Bsn* only from telencephalic pyramidal cells of the *Emx1* lineage. Bsn^Emx1^ mice do not express severe epileptic seizures (Annamneedi et al., 2018; Blondiaux et al., 2023). In the Bsn^ΔEx4/5^ mice, compared to Bsn^WT^ controls, OD-plasticity was normal in juveniles, but absent in adults, while the experience-induced enhancement of the optomotor reflex was reduced in both age groups. Notably, OD-plasticity was also absent in adult Bsn^Emx1^ mice, but spatial vision and its experience-induced changes were indistinguishable from Bsn^WT^ controls. Further, while MD induced upscaling of postsynaptic AMPA currents and enhanced surface expression of AMPARs, presynaptic upscaling was completely absent in cultured hippocampal and cortical neurons lacking *Bsn*. Finally, while MD induced profound increases in SYN phosphorylation in contralateral V1 of adult WT mice, MD-induced changes in SYN phosphorylation were substantially abolished in Bsn^Emx1^ mice. These data clearly show that presynaptic BSN is essential for adult mouse OD-plasticity, possibly mediated by experience-induced modulation of SYN phosphorylation.

## Material and Methods

### Animals

All mouse strains used in this study were characterized in previous studies. **Bsn^ΔEx4/5^** animals carry *Bsn^tm1Gund^* allele, in which the exons 4 and 5 of *Bsn* coding the central part of the protein were replaced by a lacZ/neomycin cassette resulting in the expression of a residual dysfunctional protein (Altrock et al., 2003). Mice were bred on a mixed genetic background of C57BL/6J and 129/SvEmsJ strains, which is controlled by using sustained C57BL/6J-backcrossed and 129 inbred mice to breed the heterozygous parents. In experiments, mice homozygote for *Bsn^tm1Gund^* allele were used as KO and their wildtype littermates were used as controls (Bsn^WT^). **Bsn^fl/fl^**animals carry conditional *Bsn^tm1.1Arte^* allele, in which loxP sites were inserted flanking exon 2 of the Bsn gene. Expression of cre recombinase in cells of these animals result in deletion of functional *Bsn* gene (Annamneedi et al., 2018). Conditional **Bsn^Emx1^** animals lack *Bsn* specifically at telencephalic excitatory neurons (Annamneedi et al., 2018). To generate Bsn^Emx1^ animals, conditional Bsn^fl/fl^ animals were crossed with B6.129S2-Emx1^tm1(cre)Krj^ mice expressing Cre recombinase under the control of the *Emx1* promoter in telencephalic excitatory neurons and glial cells (Gorski et al., 2002). All mice of this conditional strain used in experiments were homozygote for the *Bsn^tm1.1Arte^* allele. Bsn^Emx1^ mutants expressed the *Emx1^tm1(cre)Krj^* allele and controls (Bsn^WT^) were their littermates homozygote for the wildtype *Emx1* allele. **Bsn^GT^** mice, carrying the *BsnGT^(OST486029)Lex^* allele, were created by gene trap, characterized previously (Hallermann et al., 2010) and kept on C57BL/6N background. In experiments, homozygous Bsn^GT^ animals and their Bsn^WT^ littermates were used. Mice were housed at standard conditions (12 h light/dark cycle) with food and water provided ad libitum. Our standard cages (SCs) were 26×20×14cm large, and we housed 3-5 animals of the same litter and sex together. The cages were translucent with an open grid cover and wood chip bedding. All experimental procedures were approved by the local governments of Saxony-Anhalt, Thuringia and Lower Saxony.

### Monocular deprivation (MD)

To induce selective visual experience, the right eye of mice was sutured shut for 4 (juvenile) or 7 (young adults) days according to published protocols (Gordon and Stryker, 1996; Cang et al., 2005). In brief, mice were box-anaesthetized using 2.5 % isoflurane in O_2_:N_2_O (1:1) and received one dose of Rimadyl (5 mg/kg) bodyweight. The eyes were protected from drying using Bepanthen/Corneagel cream. 2% Lidocaine cream (2%) was applied to the eyelids, before these were trimmed and sutured shut by using 1-2 mattress stitches. Animals were checked daily to ensure that the eyes remained closed. The control group without MD was treated identically, except that no eyelids were trimmed or sutured. On the day of intrinsic signal optical imaging, the age of juvenile Bsn^WT^/Bsn^ΔEx4/5^ was 35±2/35±2 days, respectively, and the age of adult Bsn^WT^/Bsn^ΔEx4/5^ was 101±4/97±3 days, respectively. Adult Bsn^WT^/Bsn^Emx1^ animals were 94±5/90±4 days old, respectively.

### Virtual-reality optomotor setup

Both the spatial frequency and the contrast sensitivity thresholds of the optomotor reflex of all mice (animals with and without MD) were measured using the optomotor system (Prusky et al., 2004; Goetze et al., 2010). Baseline measurement was performed for both eyes to ensure balanced vision in both eyes before MD. Then MD was performed, and daily measurements continued for the left eye (open eye in MD group) for the following 4/7 (juvenile/young adults) days.

### Optical imaging of intrinsic signals and visual stimuli

After completion of the optomotor tests, visual cortical responses were recorded and analyzed as described previously (Cang et al., 2005; Goetze et al., 2010; Greifzu et al., 2011). Briefly, mice were anesthetized with 2% halothane in O_2_:N_2_O (1:1) and injected with atropine (0.3mg/mouse s.c.; Franz Köhler), dexamethasone (0.2mg/mouse s.c.; Ratiopharm), and chlorprothixene (0.2mg/mouse i.m.; Sigma). After placing animals in a stereotaxic frame, anesthesia was maintained with 0.8% halothane in a 1:1 mixture of O_2_:N_2_O. For mice with MD, the right eye was reopened at the beginning of the surgery. V1 intrinsic signals were recorded from the left hemisphere, contralateral to the deprived eye. V1-responses were recorded through the skull using the “Fourier”-imaging method of Kalatsky and Stryker (2003) and optimized for the assessment of OD-plasticity (Cang et al., 2005). V1-signals were visualized with a CCD-camera (Dalsa 1M30) using a 135×50 mm tandem lens configuration (Nikon), with red illumination light (610±10 nm). Active brain regions absorb more of the red light and appear darker in the images. Frames were acquired at a rate of 30Hz, temporally binned to 7.5Hz, and stored as 512×512 pixel images after spatial binning of the camera image. Visual stimuli were presented on a high refresh rate monitor (Benq BL240) positioned 25 cm from the eyes. During recordings, paraffin oil was applied to each eye to prevent the eyes from drying, and one eye was covered using aluminum foil for recording responses to contralateral (C) and ipsilateral (I) eye stimulation, respectively. Stimuli consisted of drifting white horizontal bars limited to cover the contralateral binocular visual field (2° wide; −5 to +15° of the visual field) as described previously (Greifzu et al., 2011). The amplitude component of the optical signal represents the intensity of neuronal activation (expressed as fractional change in reflectance ×10^-4),^ which was used for ODI calculation. The ocular dominance index (ODI) was computed as: (C−I)/(C+I), with C and I representing the response magnitudes of each pixel to visual stimulation of the contralateral and ipsilateral eye. The ODI ranges from −1 to +1, with negative values representing ipsilateral and positive values representing contralateral dominance. The quality of the retinotopic maps was assessed by the calculation described previously on contralateral eye full field maps (Cang et al., 2005). For each of the pixels in the retinotopic map, the difference between its visual field position and the mean position of its surrounding 24 pixels was calculated. For maps of high quality, the position differences are quite small because of smooth progression. The standard deviation of the position difference was then used as an index of the quality of retinotopic maps, with a small standard deviation, which corresponds to low map scatter, indicating high map quality and high values, corresponding to high map scatter, indicating low map quality.

### Primary neuronal hippocampal and cortical cultures preparation and treatments

Primary hippocampal and cortical cultures from newborn Bsn^GT^ and Bsn^Emx1^ animals and their wild-type siblings were prepared as described previously and plated in densities of 35000 and 50000, respectively, cells per coverslip (18 mm diameter) (Davydova et al., 2014). Neurons from Bsn^fl/fl^ mice were prepared as described in (Anni et al., 2021) and plated at a density of 150,000 per coverslip (18 mm diameter). Cells cultured for 2 days in vitro (DIV) were transduced with lentivirus expressing cre or control construct for cre-mediated excision of *Bsn* exon 2 leading to *Bsn* gene inactivation. For Homeostatic plasticity experiments, 18-19 DIV cells were treated for 48 hours with 50 µM of (R)-2-amino-5-phosphonopentanoic acid (APV, CAS #79055-68-8) (Tocris Bioscience) and 10 µM of 6-cyano-7-nitroquinoxaline-2,3-dione disodium (CNQX CAS #479347-85-8) (Tocris Bioscience) to induce network activity silencing. DMSO (# A994.2, Roth) was used as a vehicle control. The cells were maintained at 37°C in a humidified incubator containing 5% CO2. All chemicals used for neuronal cultures were obtained from Thermo Fisher Scientific, unless indicated otherwise.

### Electrophysiology

Whole-cell patch-clamp recordings were made at room temperature from Bsn^WT^ and Bsn^GT^ mouse hippocampal cultures (18-21 DIV) while the recording chamber was continuously perfused with extracellular solution. To chronically silence neuronal activity, we treated the neurons with APV (50 µM) and CNQX (10 µM) for 48 hr prior to the day of recording. mEPSC (miniature excitatory postsynaptic currents) recordings were performed in voltage clamp mode using borosilicate glass pipette electrodes (3-6 MΩ). Glass pipette electrodes filled with an internal solution (130 mM K-gluconate, 10 mM HEPES, 5 mM KCl, 0.1 mM EGTA, 1 mM CaCl_2_, 2 mM MgCl_2_, 4 mM Na_2_-ATP, and 0.3 mM Na-GTP; pH 7.2-7.3 adjusted with KOH) and a perfusion of extracellular solution (145 mM NaCl, 5 mM KCl, 10 mM Glucose, 10 mM HEPES, 2 mM CaCl_2_, 2 mM MgCl_2_; pH 7.4 adjusted with NaOH; 300 mOsmol) containing NMDAR blocker APV (50 µM), GABA_A_R blocker bicuculline (10 µM) and TTX (1 µM) were used to measure the AMPAR mediated mEPSCs. Neurons were held at −70 mV and the mEPSCs were recorded after a stable whole-cell voltage-clamp configuration was established, and the recording was performed for 10 min (10 sweeps). Raw data were amplified using HEKA EPC10 amplifier, and the data were acquired using PatchMaster v.2.11 software (HEKA). The sampling rate for all the recordings was 10 kHz. MiniAnalysis program version 6.0.7 (Synaptosoft) was used for the analysis of mEPSC data. 200 events from each neuron were used for the statistical analysis of mEPSC rise time (10-90%), decay time (90-37%), half width, inter-event intervals and amplitudes.

### Antibodies

The following primary antibodies were used for immunocytochemistry (ICC) in the concentrations indicated: a rabbit antibody against the SYT1 luminal domain, labeled with Oyster 550 (live staining, 1:70, #105103C3, Synaptic System, Göttingen, Germany), against SHANK2 (ICC 1:500, #162 202, Synaptic System). Mouse antibody against VGLUT1 (ICC 1:1000, MAB5502 Millipore), against pan GRIA Oyster 550-labeled (live staining 1:500, 181411C3, Synaptic System). Guinea pig antibody against SYN1/2 (ICC 1:1000, # 106004, Synaptic System). Fluorescently labeled secondary antibodies used for ICC were donkey anti-mouse Alexa 488 (ICC 1:2000, # A21202, Thermo Fisher Scientific) and Cy5-donkey anti-guinea pig (1:1000, # 706-175-148, Dianova/Jackson ImmunoResearch Labs). Primary antibodies and their respective dilutions that were use for quantitative western blots (WB) are: rabbit antibody against pS9SYN: 1:500; #NB300-180 (Novusbio), pS62SYN: 1:2000; #PA5-38336 (ThermoFisher), pS549SYN: 1:500; #NB300-744 (Novusbio), pS553SYN: 1:10000; #Ab32532 (Abcam, pS605SYN: 1:10000; #PA5-38528 (ThermoFisher) together with guinea-pig antibody against SYN1/2: 1:2000; #106 004 (Synaptic System). For fluorescent detection of WB, IRDye 800CW donkey anti-guinea pig (#926-32411) and IRDye 680RD donkey anti-rabbit (# 926-6807) from Li-COR were used.

### Immunostaining and image analysis

Immunostainings of neurons were done as described earlier (Lazarevic et al., 2011). For quantitative assessment, all coverslips compared in one experiment were processed in parallel using identical antibodies, solutions, and other reagents. For the live staining with SYT1 luminal domain antibody (SYT1Ab uptake), neurons were washed two times with fresh Tyrode’s buffer containing (119 mM NaCl, 2.5 mM KCl, 2 mM CaCl_2_, 2 mM MgCl_2_, 30 mM glucose, and 25 mM HEPES, pH 7.4) and then incubated with Oyster 550-labeled SYT1 antibody (Synaptic Systems) either for 20 minutes at 37°C diluted in the above buffer to monitor endogenous network activity-induced uptake or for 4 minutes in high K+ Tyrode’s buffer containing 71.5 mM NaCl and 50 mM KCl to assess evoked uptake. Afterward, in order to prevent unspecific labeling, cells were washed two times with physiological Tyrode’s solution prior fixation. Finally, the neurons were fixed with 4 % PFA, permeabilized and staining with antibodies against VGLUT1 to label excitatory presynapses. For surface staining of the AMPA receptor, neurons were incubated with an antibody recognizing the extracellular domain of GRIA1-4 (pan GRIA) in neuronal cell media for 30 min at 4°C to block membrane trafficking. After a brief wash with Tyrode’s buffer containing 1% BSA (Bovine serum albumin), the secondary antibody was applied in cell media at 4 °C for 10 minutes. Afterward, the cells were rewashed with Tyrodes buffer + 1% BSA and processed for immunostaining as described above.

Images of stainings were acquired on a Zeiss Axio Imager A2 microscope with Cool Snap EZ camera (Visitron Systems) controlled by VisiView (Visitron Systems GmbH) software. For quantifications, settings of camera or photomultiplier were applied identically to all coverslips quantified in one experiment. For each IF quantification in each experiment, images from at least two different coverslips were acquired and quantified to avoid effects given by experimental variance. Unspecific background was removed using threshold subtraction in ImageJ software (NIH, https://imagej.net/ij/). For analysis of SYT1Ab uptake, synaptic puncta were defined semiautomatically by setting rectangular regions of interest (ROI) with dimensions of about 0.8X0.8um around local intensity maxima in the channel with staining for synaptic marker SYT1 and VGLUT1 using OpenView software (written and kindly provided by N.E. Ziv (Tsuriel et al., 2006). Surface AMPA receptor staining experiments were analyzed analogously with SYN1 and SHANK2 as synaptic markers. Mean IF intensities were measured in synaptic ROIs in all corresponding channels using the same software and normalized to the mean IF intensities of the control group for each of the experiments.

### Lentivirus particle production

The lentiviral vector for expression of ratio:sypHy (Rose et al., 2013) and active and inactive cre recombinase and their production was described (Kaeser et al., 2011; Lazarevic et al., 2017; Anni et al., 2021).

### Synapto-pHluorin (sypHy) Imaging

To quantify the size of readily releasable pool (RRP) and total recycling pool (TRP) imaging of sypHy fluorescence was carried out in mature hippocampal neurons (DIV19-21) transduced with the lentiviral particles on day 4. Coverslips with neurons expressing sypHy were placed in a field stimulation chamber (RC-49MFSH; Warner instruments) and imaged at RT on an inverted microscope (Zeiss Axio Observer.A1) equipped with an EMCCD camera (Evolve 512; delta Photometrics) controlled by VisiView (Visitron Systems GmbH). A 63x oil immersion objective NA 1.4 and GFP/mCherry single band exciters ET filter set (exciter 470/40, exciter 572/35, emitter 59022m, dichroic 59022BS) were used. Transduced neurons were identified by their RFP expression at the beginning of the experiment. Neurons were stimulated in the presence of 50 µM APV and 25 µM CNQX to avoid recurrent activity and 1µM of Bafilomycin A, a reversible inhibitor of the vesicular proton pump (V-type ATPase), to prevent the vesicle re-acidification of retrieved vesicles (Sankaranarayanan and Ryan, 2001). After 5s of baseline acquisition (F_0_), neurons were stimulated via field electrodes to elicit 40 pulses at 20Hz ensuring the release of the vesicles from RRP, followed by 2 min recovery time. After that the TRP was released by application of 900AP at 20Hz and at the end a pulse of 60 mM NH_4_Cl (F_max_) containing solution was applied (Burrone et al., 2006) to estimate the total sypHy-expressing vesicle pool. Electrical stimulation was generated using an isolator unit (World Precision Instruments) controlled by a pulse generator (Master-8). Imaging was performed at a frequency of 12 Hz during 30 sec for the RRP and during 60 sec for the TRP. Synaptic boutons that responded to the stimulation were selected by subtracting the first 10 frames from the baseline from the 10 frames directly after the stimulus. Only the boutons showing a difference between the TRP fluorescence and NH_4_Cl-evoked fluorescence higher than 20%, were considered for the analyzes. The mean IF intensities were measured in the circular regions of interest (ROIs with a diameter of 8 × 8 pixel) placed over each responding bouton using Time Series Analyzer V2.0 plugin in ImageJ (NIH, https://imagej.net/ij/). Mean IF values from between 80 and 150 ROIs/visual field per cell were analyzed and plotted after bleaching correction using GraphPad Software. The relative sizes of the RRP and the RP were expressed as fractions of the total sypHy-expressing pool detected after addition of NH_4_Cl. RRP and TRP were quantified by averaging the mean of 100 values per each cell (representative of the frames 197–297 corresponding to time points 15–24 s and 600-700 corresponding to time points 48.8 - 56.08 s, respectively, on the XY graph).

### Quantitative western blot and analyses

Ipsilateral and contralateral sides of V1 from MD and mock treated animals (sham condition) control (Bsn^WT^) or Bsn ^Emx1^ mice were collected and homogenized using a teflon-homogenizer in the presence of a homogenization buffer (2M sucrose, 500mM HEPES, Complete Protease inhibitor Cocktail (Roche #04693116001), PhosSTOP^TM^ (Roche #04906837001). Lysates were collected and then centrifuged at 13,000 RPM, 4 °C for 10 min. The supernatant was collected and protein concentration was determined using standard BCA assay. 10µg of proteins were separated in 7.5% TGX^TM^ (Bio-Rad #4568023) stain free gels fitted in a Bio-Rad 2-D electrophoresis chamber (Bio-Rad #1688005) and then blotted onto Bio-Rad low fluorescence PVDF membranes under semi-dry transfer conditions in Bio-Rad Turbotransfer^TM^ system. Each sample was loaded twice into the gels. Fluorescent immunodetection was carried out in a LI-COR Odyssey^TM^ infrared scanner (at the signal emission spectrum of 680 nm or 800 nm respectively). Immunofluorescence (IF) signals corresponding to phosphorylation states of SYN phospho-species pS9SYN, pS62pSYN, pS549SYN, pS553SYN, pS605 SYN and total SYN were quantified using the detection and quantification software Image Studio V5.2 (LI-COR Biosciences). The IF of each detected SYN isoforms (Ia, Ib, IIa, IIIa and IIb) for respective SYN phospho-species from either the contralateral or ipsilateral side of V1 were first normalized to the TCE staining of total protein in each slot, which served as a loading control. These normalized IF values of SYN isoforms from each immunodetected SYN phospho-species were then expressed as values relative to the averaged IF value detected in the contralateral and ipsilateral side of V1 of the respective animal (relative contra IF = IFcontra /0.5*(IFcontra + IFipsi) and relative ipsi IF = IFipsi / 0.5*(IFcontra + IFipsi). Each sample was loaded twice and an average was made between the two quantified IFs or final values. These values were then plotted onto graphs using Graphpad Prism 9.5.1. Samples from 3 animals per condition were utilized in the final analysis. All data points for quantitative analyses are represented as mean ± SEM in text. Statistical analyses were performed using Multiple unpaired t-test with Holm-Sidak post-test, following statistical assumption of normality distribution.

### Statistical analysis

All results of quantitative analyses are given as means ± standard errors of the mean (SEM) in figures and data tables. Statistical analyses were performed with Prism 11 (in vitro experiments) or 8 (in vivo experiments) software (GraphPad Software, Inc.). Upon assessment of normal distribution, inter-group comparisons between two groups were done by Student’s t-test and for 3 groups and more using one, two or three-way analysis of variance (ANOVA), as detailed in the text. In analyses in which a within-subject factor was present (i.e., eye, age or spatial frequency), ANOVA with repeated measurements (rm) was performed. Post hoc multiple comparison tests for *in vivo* data in figures 1, 2 and S1 were corrected for false discovery rate (FDR) using the method by Benjamini and Hochberg unless stated otherwise. Statistical analyses of quantitative WBs were performed using Multiple unpaired t-test with Holm-Šídák post-test, following statistical assumption of normality distribution. Statistical significance and information about group character and size are specified in the figure legend for each experiment and in Table 1.

**Figure 1:**
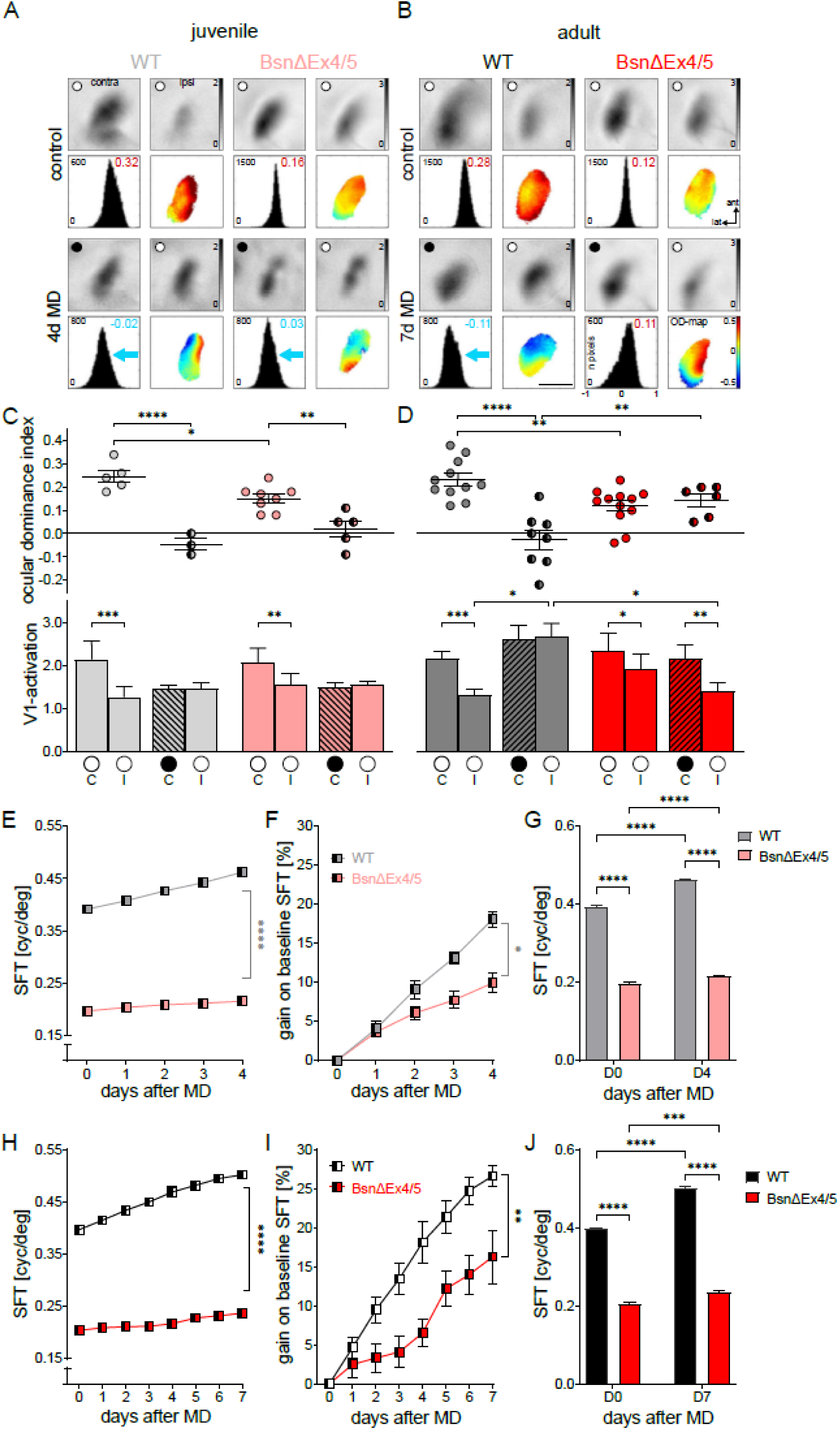
Adult Bsn^ΔEx4-5^ mice failed to display ocular dominance plasticity in primary visual cortex (V1) and both juvenile and adult Bsn^ΔEx4-5^ mice had reduced spatial vision and compromised experience-induced enhancements of the optomotor reflex. (A, B) Optically recorded activation in the binocular region of mouse (V1) induced by visual stimulation of the contralateral (contra) and ipsilateral (ipsi) eye in juvenile (A) and adult (B) Bsn^WT^ and Bsn^ΔEx4-5^ mice. Upper/lower 2 rows: V1-activity maps and their quantification in control/MD-mice. Open/closed eyes (after MD) are indicated by a white/black circle, respectively. Grayscale-coded V1-response magnitude maps after contra- and ipsilateral eye visual stimulation (top row), histogram of OD-scores including ocular dominance index (ODI) and 2-dimensional OD-maps (lower row) are illustrated. In all mice without MD (control), V1-activity patches evoked by stimulation of the contralateral eye were darker than activity patches after ipsilateral eye stimulation, the ODI was positive, and warm colors prevailed in OD-maps, indicating contralateral dominance. (C, D) Quantification of V1-activity maps in juvenile and adult Bsn^WT^ (grey/black) and Bsn^ΔEx4-5^ (rose/red) mice. Both average ODI (top row) and average V1-activation after ipsi (I)/contralateral (C) eye stimulation (lower row) are displayed. ODI: symbols represent individual values; means are marked by horizontal lines. In juvenile animals of both genotypes, 4 days of MD induced an OD-shift towards the open (experienced) eye: ODIs after MD were significantly lower as without MD. In contrast, in adult animals, 7 days of MD induced an OD-shift only in Bsn^WT^- but not in Bsn^ΔEx4-5^ mice. After MD, V1-activation (hatched bars) was similar after contra- and ipsilateral eye stimulation in juveniles of both genotypes and in adult wildtypes. In juveniles, the OD-shift was primarily mediated by a reduction in V1-activation via contralateral (closed) eye stimulation. In contrast, in adult Bsn^WT^ mice, the OD-shift after MD was primarily mediated by an increase of open eye V1-activation, which was absent in adult Bsn^ΔEx4-5^ mice. (E-J) Optomotor measurements of the spatial frequency threshold (SFT) of the open eye before and during MD in juvenile (E-G) and adult (H-J) Bsn^WT^ and Bsn^ΔEx4-5^ mice. While baseline SFT of both juvenile and adult Bsn^ΔEx4-5^ mice was much lower than SFT of Bsn^WT^ mice, both genotypes and age groups increased SFT after MD. However, gain on baseline SFT after 4/7 days of MD was lower in Bsn^ΔEx4-5^ compared to Bsn^WT^-mice. 2-way ANOVA (C, D - top row and G, J) and 3-way ANOVA (C, D - bottom row and E, F, H, I) was used to test main effects and interactions. Post hoc multiple comparisons were corrected for false discovery rate (C, D) or Sidak (E-J) *p<0.05; **p<0.01; ***p<0.001; ****p<0.0001

**Figure 2:**
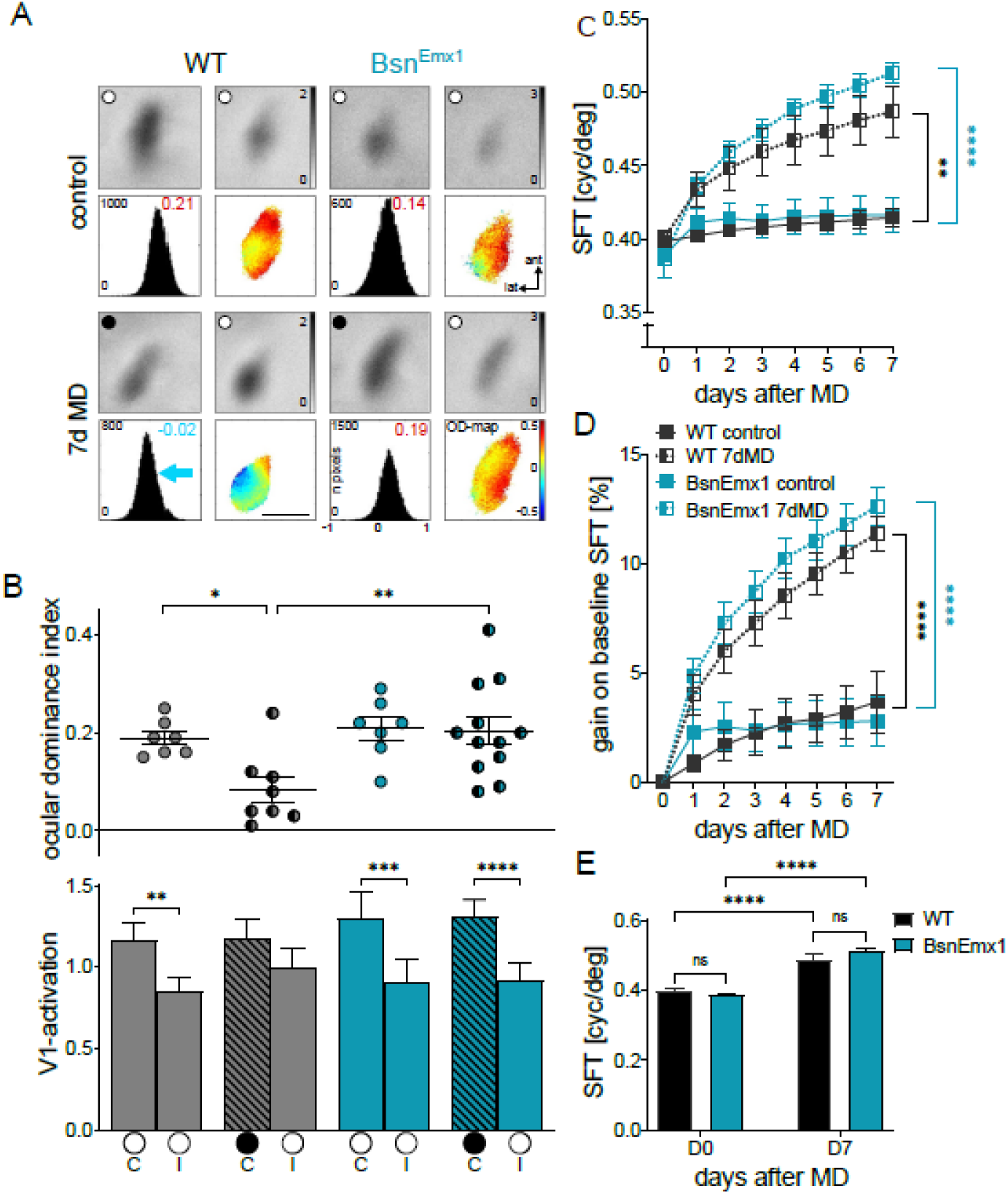
Adult Bsn^Emx1^ mice failed to display ocular dominance plasticity in V1. Optomotor measurements Data display and abbreviations as in Fig. 1. (A) V1-activity maps of adult (>P80) Bsn^WT^ and Bsn^Emx1^ mice with and without MD. (B) Quantification of V1-activation. While Bsn^WT^ mice show an OD-shift towards the open eye after seven days of MD, OD-plasticity is absent in adult Bsn^Emx1^ mice, and V1-activation remains dominated by contralateral eye input. (C-E) Baseline SFT of Bsn^WT^ and Bsn^Emx1^ mice is similar, and SFTs get enhanced after MD in both Bsn^WT^ and Bsn^Emx1^ mice. (C) Spatial frequency threshold of the optokinetic response in cycles per degree (cyc/deg) plotted against days after MD (or control). (D) Gain on baseline SFT in percent (%) plotted against days after MD (or control). (E) Comparison of the SFT on day 0 and after 7 days of MD. 2-way ANOVA (B top, E) and 3-way ANOVA (B bottom, C, D) was used to test main effects and interactions. Post hoc multiple comparisons were corrected for false discovery rate (B) or Sidak (C-E); *p<0.05; **p<0.01; ***p<0.001; ****p<0.0001.

## Results

### OD-plasticity is impaired in adult, but not in young mice with a constitutive deletion of *Bsn*

One of the most important questions of the present study was whether the impairment of synaptic function in *Bsn* mutant mice would interfere with visual cortical plasticity *in vivo* (Goetze et al., 2010; Montenegro-Venegas et al., 2022). We therefore used the established paradigm of OD-plasticity after monocular deprivation (MD) in a mouse model with globally disrupted *Bsn* expression (Bsn^ΔEx4-5^). Using optical imaging of intrinsic signals, we compared the response amplitudes in the binocular region of primary visual cortex (V1) after stimulating the ipsi- or contralateral eye with moving horizontal bars (Cang et al., 2005; Goetze et al., 2010). We imaged V1-activity maps of both juvenile (postnatal day (P) 28-35) and young adult (P83-105) Bsn^ΔEx4-5^ mice and their wildtype littermates (Bsn^WT^) without (noMD) and after 4 (juvenile) or 7 (adult) days of MD.

In Bsn^WT^ controls and Bsn^ΔEx4-5^ mice without MD, visual stimulation of the contralateral eye induced stronger stimulus-evoked V1-activation than stimulation of the ipsilateral eye, demonstrating that the contralateral eye dominates activity in binocular V1 in both genotypes (Figure 1A, B top row, Table 1). To quantify the relative strength of V1-activation after left and right eye stimulation, we computed an OD-index (ODI), and this index is color-coded in the two-dimensional OD-map, with warm colors and positive values indicating contralateral dominance, and cold colors / negative values indicating ipsilateral dominance (Figure 1A, B, second row). Notably, both non-deprived juvenile and adult Bsn^ΔEx4-5^ mice had a lower average ODI of 0.15±0.02/0.12±0.02 (juvenile/adult) compared to juvenile/adult 0.25±0.03/0.24±0.02 Bsn^WT^ mice (p=0.017/0.004; Figure 1A-D). Thus, while the contralateral eye still dominates binocular V1-activity after systemic *Bsn* deletion, in non-deprived animals of both age groups, this dominance was significantly reduced compared to Bsn^WT^ mice.

After establishing baseline V1-activity after ipsi- and contralateral eye stimulation, we next analyzed the effects of 4 or 7 days of MD in young and adult Bsn^ΔEx4-5^ animals. In juvenile mice of both genotypes, V1-responses induced via either eye became more similar resulting in a lowered OD-index (Figure 1A, third and fourth row), with Bsn^WT^ and Bsn^ΔEx4-5^ showing average ODIs of 0.05±0.03 and 0.02±0.03, respectively (Figure 1C, Table 1; Bsn^WT^/Bsn^ΔEx4-5^ p-value <0.0001/<0.01). In both genotypes, OD-plasticity was primarily mediated by decreased V1-responses to visual stimulation of the (previously) deprived, contralateral eye, resulting in rather similar V1-activation by both eyes (Figure 1A, third row, and 1C). Both the magnitude of the observed OD-shifts and the V1-activation changes were in line with previous studies done in C57Bl/6J mice of a similar age range (Cang et al., 2005; Lehmann and Löwel, 2008).

In contrast to juvenile mice, adult Bsn^ΔEx4-5^ mice failed to express OD-plasticity (Fig. 1B): While in adult Bsn^WT^-mice, V1-activation via either eye was comparable after 7 days of MD, in Bsn^ΔEx4-5^ animals, the deprived (contralateral) eye continued to dominate V1-activity (Figure 1B, third row). Quantitative analyses confirmed that OD-plasticity was absent in Bsn^ΔEx4-5^ mice. In adult Bsn^WT^ mice, the ODI shifted from 0.23±0.03 without MD to −0.03±0.04 after 7 days of MD (Figure 1D, Table 1; p<0.0001). In contrast, in adult Bsn^ΔEx4-5^ mice, ODIs before and after MD were rather similar: 0.12±0.02 (noMD) and 0.14±0.03 (MD), i.e. there was no OD-shift after MD (Figure 1D, Table 1; p>0.05). Thus, ODIs after 7d MD were significantly different between both genotypes (Figure 1D, p=0.004). In Bsn^WT^ mice, the OD-shift was clearly mediated by an increased V1-activation via the open (ipsilateral) eye (Figure 1D; V1-activation, noMD/7d MD: 1.44±0.18/2.66±0.36, p<0.05). In contrast, in Bsn^ΔEx4-5^ mice, open eye mediated V1-activation did not increase after MD (Figure 1D; control/7d MD: 1.91±0.37/1.39±0.22, p>0.05; Table 1). V1-activation of the closed (contralateral) eye remained unchanged irrespective of genotype.

Overall, our results show that both juvenile and adult Bsn^ΔEx4-5^ mice have a significantly lower ODI compared to Bsn^WT^-animals (without MD), indicating a reduced contralateral dominance in binocular V1. This did not interfere with OD-plasticity in juvenile Bsn^ΔEx4-5^ mice, which was primarily mediated by a decrease in V1-activation via the deprived eye, similar to Bsn^WT^ animals. In contrast, OD-plasticity was absent in adult mice with a constitutive deletion of functional *Bsn*: specifically, the MD-induced potentiation of open (ipsilateral) eye V1-activation normally observed after 7 days of MD in Bsn^WT^ mice, did not occur.

## Impaired experience-induced enhancement of the spatial frequency threshold of the optokinetic

### reflex in Bsn^ΔEx4-5^ mice

We previously showed that the spatial frequency threshold (SFT) of the optokinetic response of Bsn^ΔEx4-5^ mice to moving vertical sine wave gratings is reduced by about 50% (Goetze et al., 2010). Here we asked whether the experience-induced SFT-enhancement known to occur after MD (Prusky et al., 2006), is also impaired.

In line with published observations (Goetze et al., 2010), both juvenile and adult Bsn^ΔEx4-5^ mice had a drastically reduced SFT of 0.20±0.003 (juvenile)/0.20±0.004 cyc/deg (adult) compared to their wildtype littermates with 0.39±0.004 (juvenile)/0.40±0.002 cyc/deg (adult) (Figure 1E-J). Nevertheless, following MD, the SFT of the open eye increased in all groups (Figure 1E-J, Table 1). In juvenile/adult Bsn^WT^ animals, the SFT increased to 0.46±0.003/0.50±0.004 cyc/deg on day 4/7 after MD (D0 vs D4/7: p=0.0004/p<0.0001) (Figure 1E). In contrast, in juvenile/adult Bsn^ΔEx4-5^ mice, the SFT increased to only 0.22±0.003/0.24±0.003 cyc/deg (D0 vs D4/7: p=0.0001/ p<0.0001) (Fig. 1H).

Since Bsn^ΔEx4-5^ mice started from a substantially lower SFT, we also compared the relative experience-induced increase of SFT after MD (gain on baseline) between genotypes by calculating the percent increase in SFT relative to the starting value before MD for each age/genotype (Figure 1F and I). In juvenile/adult Bsn^WT^ mice, the gain on baseline after MD was 18±1/27±1 % and thus significantly higher than the 9±1/16±3 % of the age-matched Bsn^ΔEx4-5^ mice (RM 2-way ANOVA – effect of genotype; juveniles/adults: p=0.011/0.003). Thus, the constitutive deletion of functional *Bsn* not only reduces baseline spatial vision as shown previously, but it also dramatically reduces the experience-induced enhancement of the optomotor reflex of the open eye after MD.

### MD-induced enhancement of the contrast sensitivity is impaired in Bsn^ΔEx4-5^ mice

MD not only induces an increase in the SFT, it also increases the contrast sensitivity of the optokinetic response of the open eye (Prusky et al., 2006). To analyze the consequences of the constitutive deletion of functional *Bsn* on this parameter, we performed daily measurements of contrast sensitivity at six different spatial frequencies in both non-deprived and MD-animals of both genotypes and age groups. The highest contrast sensitivity was measured at 0.064 cyc/deg for all animals. However, and consistent with previous findings (Goetze et al., 2010), baseline contrast sensitivity of Bsn^ΔEx4-5^ mice was dramatically reduced at all spatial frequencies and only amounted to about 20% to 50% of the control littermate’s values (Supplementary figure 1A-F, Table S1), i.e. Bsn^ΔEx4-5^ mice needed much higher contrasts to elicit an optomotor response. Following MD, contrast sensitivity of the open eye significantly increased across days for all spatial frequencies in both age and genotype groups (Supplementary figure 1A-F; Table S1). At the spatial frequency of 0.064 cyc/deg, juvenile Bsn^WT^ mice increased their contrast sensitivity from 15.9±0.6 (corresponding to 6.3% contrast) before deprivation to 22.5±0.9 (4.4% contrast) on day 4 after MD (Supplementary figure 1A, E). However, contrast sensitivity of juvenile Bsn^ΔEx4-5^ mice only increased from 2.7±0.1 (36.7% contrast) before MD to 3.1±0.1 (32.2% contrast) on day 4 after MD (Supplementary figure 1C, E). The experience-induced increase in contrast sensitivity thresholds in juvenile Bsn^ΔEx4-5^ mice was thus substantially impaired, being about three-fold lower compared to Bsn^WT^ mice (Supplementary figure 1E, Table S1, Bsn^WT^/ Bsn^ΔEx4-5^: 43±8%/14±2%, mixed-effects model, effect of genotype p=0.0014).

As expected for adult Bsn^WT^ animals, at 0.064 cyc/deg, contrast sensitivity increased from 15.6±1.0 (6.4% contrast) before MD to 28.0±3.3 (3.6% contrast) on day 7 after MD (Supplementary figure 1B, F). In contrast, in adult Bsn^ΔEx4-5^ mice, contrast sensitivity only increased from 2.7±0.1 (37% contrast) before MD to 3.0±0.1 (33.2% contrast) on day 7 after MD (Supplementary figure 1D, F). The relative increase in contrast sensitivity after MD was 84±23% in adult Bsn^WT^ animals, compared to only 14±3% in adult Bsn^ΔEx4-5^ mice, which again was significantly lower (Supplementary figure 1F, Table S1, mixed-effects model, effect of genotype: p=0.049).

Together, our detailed optomotor reflex measurements confirm substantial impairments in baseline contrast sensitivity of Bsn^ΔEx4-5^ mice compared to their Bsn^WT^ littermates, with Bsn^ΔEx4-5^ mice requiring visual stimuli with an about 6x higher contrast to induce an optokinetic reflex. In addition, the experience-induced improvement of contrast sensitivity thresholds after MD was substantially impaired in Bsn^ΔEx4-5^ mice, demonstrating that constitutive deletion of functional *Bsn* severely compromises both basic spatial vision and the experience-induced reflex enhancements.

**Supplementary figure 1:**
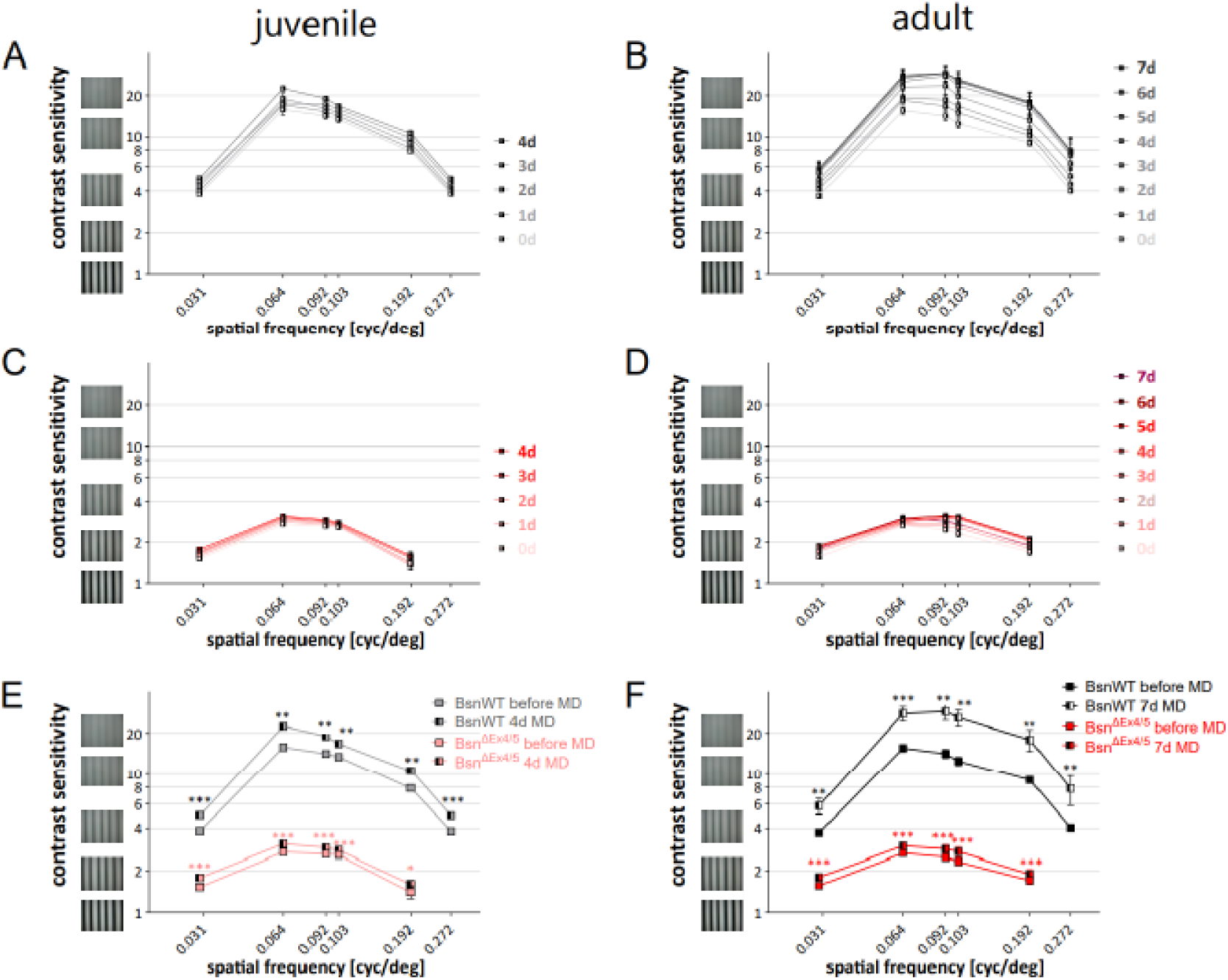
Contrast sensitivity thresholds and their experience-enabled enhancement in juvenile and adult Bsn^WT^ and Bsn^ΔEx4,5^ mice. (A-D) The optomotor reflex of the open eye was tested at a range of spatial frequencies and contrasts: contrast sensitivity thresholds are plotted as a function of the spatial frequency of the test gratings in cycles per degree (cyc/deg) for juvenile (A, C) and adult (B, D) Bsn^WT^ (grey/black lines, A, B) and Bsn^ΔEx4-5^ mice (rose/red, C, D) for each day after MD. During MD, all groups showed significant enhancements of their contrast sensitivity. (E, F) Comparison of contrast sensitivity thresholds on the first and last day of MD for juvenile (E) and adult (F) mice. In both age groups, and before and after MD, contrast sensitivity values of Bsn^ΔEx4-5^ mice were much lower than values of Bsn^WT^ animals. Mixed effects analysis was used to test main effects and interactions. Post hoc multiple comparisons were corrected for false discovery rate. *p<0.05; **p<0.01; ***p<0.001.

### Ocular dominance plasticity is also impaired in adult mice with a deletion of *Bsn* only in cortical excitatory neurons

Since Bsn^ΔEx4-5^ mice show epileptic seizures, which might contribute to the observed deficits in both vision and experience-dependent visual plasticity (Altrock et al., 2003; Blondiaux et al., 2023), we also analyzed adult animals of the Bsn^Emx1^ strain, in which *Bsn* expression is selectively abolished in cortical excitatory neurons, but preserved in the subcortical visual pathway (Annamneedi et al., 2018). These mice show normal explorative behavior and only sporadic epileptic seizures (Annamneedi et al., 2018; Blondiaux et al., 2023).

Intrinsic signal optical imaging of V1-activity in adult (non-deprived) Bsn^Emx1^-mice showed a clear dominance of the contralateral eye in binocular V1, i.e. significantly higher V1-activation after visual stimulation via the contralateral compared to the ipsilateral eye, as also observed in Bsn^WT^ control mice (Figure 2A upper row, Table 2, contra vs ipsi: p<0.05 in both genotypes). Accordingly, quantitative analyses revealed that ODIs of Bsn^Emx1^ mice were comparable to ODIs of Bsn^WT^ mice (Figure 2A second row, 2B; p=0.80).

After 7d MD, however, Bsn^Emx1^ mice, in contrast to the Bsn^WT^ controls, did not show OD-shifts in V1, and the deprived (contralateral) eye continued to dominate V1-activity. Hence, while in Bsn^WT^-mice, MD lowered the ODI from 0.19±0.01 (without MD) to 0.08±0.03 (after MD, p=0.023), in Bsn^Emx1^ mice, the ODI remained rather constant, with 0.21±0.02 without MD and 0.20±0.03 after MD (p=0.903) (Figure 2A third row, 2B, Table 2, contra vs ipsi, Bsn^WT^/ Bsn^Emx1^: p=0.11/<0.05).

These results document that deletion of *Bsn* only from cortical glutamatergic neurons, as in Bsn^Emx1^ mice, also prevents OD-shifts after MD in V1 of adult mice, as we had observed in Bsn^ΔEx4-5^ mice. This observation confirms the importance of the presynaptic scaffold protein BSN for OD-plasticity in adult mouse V1, and indicates that expression of *Bsn* specifically in neocortical glutamatergic neurons is required for these experience-dependent activation changes to occur.

### Normal spatial vision and experience-enabled enhancements of the optomotor reflex in mice with a conditional deletion of Bsn from forebrain neurons

Next, we tested the SFT of the optomotor response to vertical sine wave gratings in Bsn^Emx1^ and Bsn^WT^ mice. Since the optomotor reflex is largely subcortically mediated (Douglas et al., 2005) we did not expect to observe compromised basic spatial vision (in contrast to what we had measured in Bsn^ΔEx4-5^ mice). As hypothesized, baseline SFT of Bsn^Emx1^ mice was 0.39±0.00 cyc/deg, and thus identical to the 0.39±0.02 cyc/deg of control Bsn^WT^ mice (Figure 2C, E; Table 2). After MD, SFT-thresholds increased independent of genotype: in Bsn^WT^/Bsn^Emx1^ mice, SFT increased to 0.49±0.00/0.51±0.01 cyc/deg, respectively, after 7 days of MD (Figure 2C, E, Table 2). Likewise, the gain on baseline SFT revealed no significant difference between genotypes (Figure 2D; Table 2): 11.4±0.8/12.7±0.9 % for Bsn^WT^/Bsn^Emx1^ mice. Our results show that deletion of *Bsn* from neocortical glutamatergic networks does neither interfere with normal optomotor reflexes, nor their experience-induced threshold increases.

### Presynaptic, but not postsynaptic, inactivity-induced scaling is impaired upon *Bsn* deletion *in vitro*

A cellular mechanism that had been linked to OD-plasticity and which has age-specific features is inactivity-induced synaptic scaling (Sato and Stryker, 2008; Wen and Turrigiano, 2021). During synaptic scaling, neurons adjust their synaptic transmission to keep the overall firing frequency within physiologically meaningful rates (Maffei and Turrigiano, 2008; Turrigiano, 2008; Turrigiano, 2012). Both *in vivo* and *in vitro* studies showed that synaptic scaling can involve changes in the efficacy of neurotransmitter release as well as changes in numbers and properties of postsynaptic neurotransmitter receptors (Tropea et al., 2009). While *Bsn* is a major regulator of presynaptic plasticity its requirement for synaptic scaling has not yet been addressed (Altrock et al., 2003; Mendoza Schulz et al., 2014).

To close this gap, we assessed inactivity-induced synaptic scaling in cultured primary hippocampal neurons derived from a Bsn^GT^ strain, with a constitutive deletion of functional *Bsn*. This strain phenocopies the synaptic and systemic phenotypes described in the original Bsn^ΔEx4-5^ strain (Hallermann et al., 2010; Ryl et al., 2021; Montenegro-Venegas et al., 2022; Blondiaux et al., 2023). To trigger inactivity-induced synaptic upscaling in cultures grown for 18-19 days in vitro (DIV), network activity was silenced by application of glutamate receptor antagonists (50µM D-APV5 and 10µM CNQX) for 2 days. This treatment was previously shown to mediate pre-and postsynaptic homeostatic upscaling (Thiagarajan et al., 2005; Wierenga et al., 2005; Lazarevic et al., 2011). To assess synaptic scaling, we recorded miniature excitatory postsynaptic currents (mEPSCs) in a whole-cell patch-clamp configuration and analyzed their kinetics, frequency and amplitude (Fig 3A, Supplementary figure 2). Genotype and treatment had no effect on kinetics of mEPSCs (Table S2). In silenced Bsn^WT^ cells, mEPSC inter-event intervals (IEI) were reduced by about 80% and mEPSC amplitude increased by about 60% as compared to values measured in cells with basal activity (ctrl) (Fig.3 B-D, Table 3). Thus, pre- and postsynaptic upscaling was successfully induced by inactivity in our experimental setting. In neurons from Bsn^GT^ animals, the silencing of network activity induced an increase in mEPSC amplitudes indistinguishable from the effect observed in Bsn^WT^ cells (Fig. 3A, D, E). However, silencing had no effect on the mEPSC inter-event interval in Bsn^GT^ neurons indicating that *Bsn* is required for inactivity-induced presynaptic scaling (Fig 3A, B). Of note, at baseline activity, the Bsn^GT^ neurons showed normal amplitudes, but significantly larger inter-event intervals compared to controls (Fig. 3A-E). This indicates unchanged postsynaptic properties, but lower basal presynaptic strength, which is in line with the previously published data (Altrock et al., 2003; Hallermann et al., 2010; Davydova et al., 2014; Montenegro-Venegas et al., 2022). The increased mEPSC amplitudes have been linked to an increased externalization of AMPA-type glutamate receptors (GRIAs) in silenced cells (Thiagarajan et al., 2005; Wierenga et al., 2005). We therefore labeled GRIAs localized on the synaptic surface with an antibody recognizing extracellular domain of GRIA 1-4 in living cells. A sequential postfixation labeling with the presynaptic marker SYN was used to identify synaptic puncta. In line with normal inactivity-induced upscaling of mEPSC amplitude in Bsn^GT^, postsynaptic surface abundance of GRIAs increased by about 50% in silenced Bsn^WT^ and Bsn^GT^ neurons (Supplementary figure 3A,B) further confirming normal postsynaptic scaling in the absence of *Bsn*. An analogous experiment performed using cortical Bsn^KO^ neurons, generated upon expression of cre in conditional Bsn^fl/fl^ cells that express the same *Bsn^tm1.1Arte^* allele as Bsn^Emx1^ stain (Figure 3F-G). These data revealed a specific deficit in the presynaptic component of inactivity-induced synaptic scaling upon deletion of *Bsn*.

**Figure 3:**
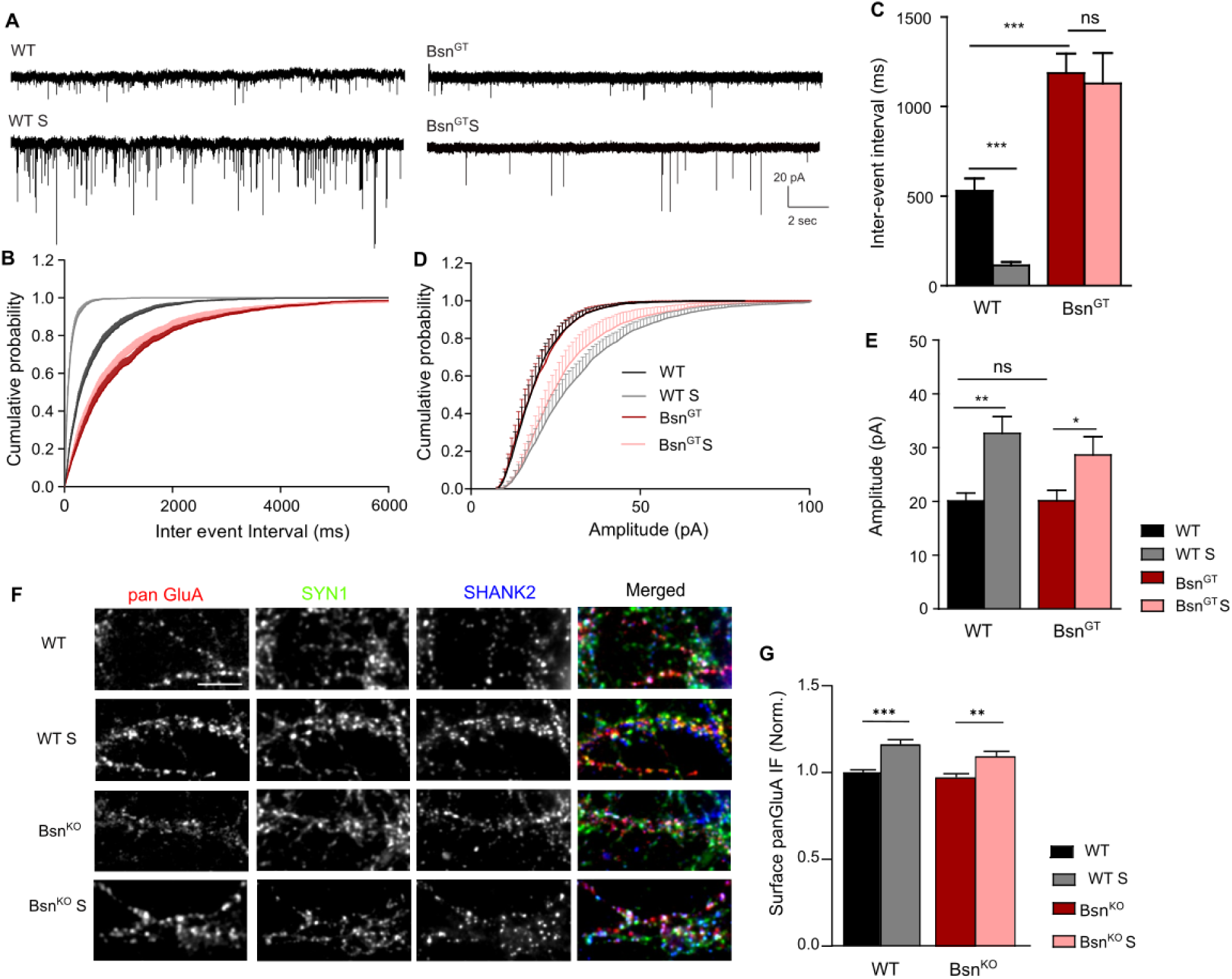
Presynaptic, but not postsynaptic inactivity-induced synaptic scaling is disrupted upon deletion of *Bsn*. (A) Representative raw traces of mEPSC recorded from control or silenced (S) neurons with gene trap-mediated *Bsn* deletion (Bsn^WT^ and Bsn^GT^). (B, D) Cumulative histograms of mEPSC inter-event intervals (IEI, B) and amplitudes (D) showing the robust increase in IEI (decreased frequency) and amplitudes after chronic silencing for 48hr in Bsn^WT^ neurons, and an increased amplitude without any significant change in IEI in Bsn^GT^ neurons. (C, E) Quantification of mEPSC IEI (C), amplitude (E) of mEPSC events recorded from control and silenced Bsn^WT^ (vehicle, n-10; S, n-12) or Bsn^GT^ (vehicle, n-10; S, n-10) neurons. Data expressed as Mean ± SEM and Significance was tested with two-way ANOVA with Tukey’s post hoc test; *p<0.05, **p<0.01, ***p<0.001, ns – not significant; n=number of cells recorded from 3 independent cultures. (F) Example images show surface population of AMPARs revealed by immunolabeling with N-terminal anti panGRIA antibody in living neurons, with cre-mediated deletion of *Bsn* kept at basal activity and upon silencing (S). Synaptic puncta were visualized by post-fixation staining with anti SYN antibody. (G) Quantification of the surface expression of glutamate receptors experiment in F. Significance was assessed with the two-way ANOVA with Sidak Post-test; **p<0.01, ***p<0.001. n=number of cells recorded from 3 independent cultures.

**Supplementary figure 2:**
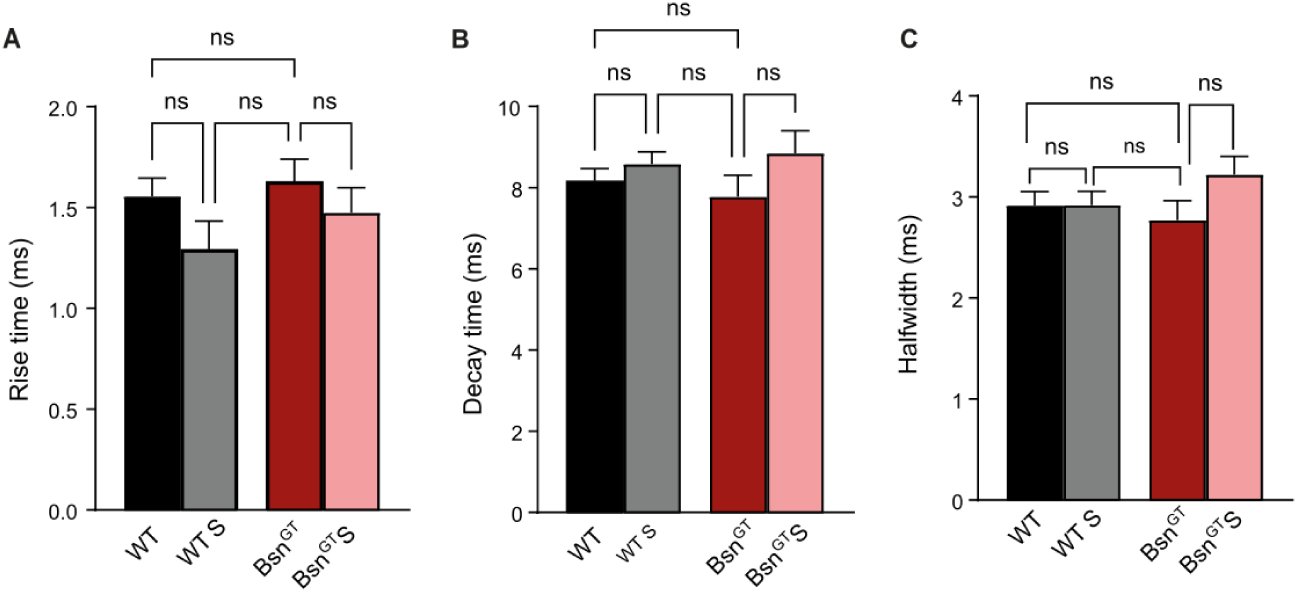
Silencing has no effect on kinetics parameters of mEPSC in control and Bsn^GT^ animals. Quantification of mEPSC kinetics parameters including rise time (A), decay time (B) and halfwidth (C) of mEPSC events recorded from treated Bsn^WT^ and Bsn^GT^ neurons kept at basal activity or silenced (S) for 48 hrs. No significant changes in the mEPSC kinetics parameters were observed neither in silenced Bsn^WT^ (vehicle, n-10; S, n-12) nor Bsn^GT^ (vehicle, n-10; S, n-10) neurons. Data expressed as Mean ± SEM and Significance was tested with two-way ANOVA with Tukey’s post hoc test; ns – not significant; The graphs show the results from 3 independent cultures.

**Supplementary figure 3:**
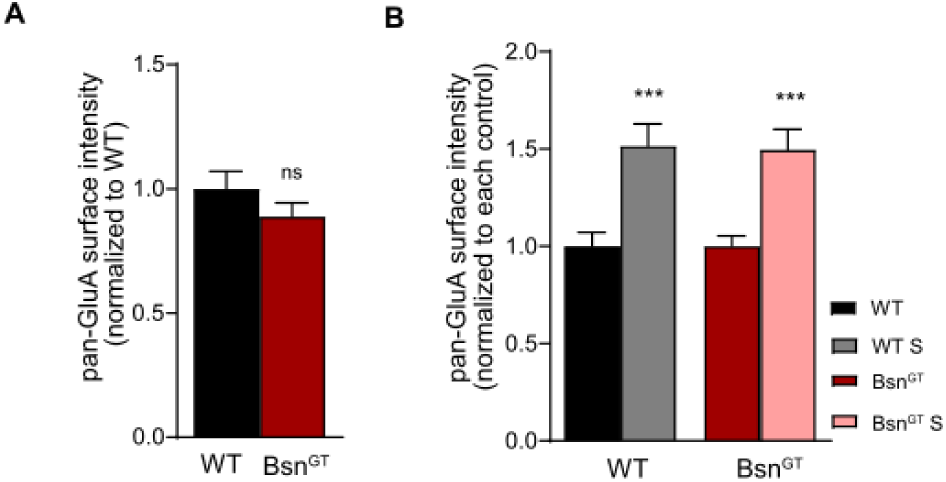
Silencing-induced increase in surface expression of AMPA receptors is identical in neurons from control and Bsn^GT^ animals. (A, B) The surface population of AMPARs was quantified by immunolabeling with an N-terminal anti-panGluA antibody in living neurons from Bsn^WT^ and Bsn^GT^ mice under basal conditions and after 48 hours of silencing. Significance was assessed with the Student’s t-test; ***p< 0.001. The graphs show results from 3 independent cultures.

### *Bsn* is necessary for inactivity-induced homeostatic synaptic scaling in cultured neurons

In presynapses, only a fraction of neurotransmitter filled synaptic vesicles (SVs) contribute to neurotransmission and a considerable number of SVs do not undergo exocytosis even after prolonged trains of stimulation (Rizzoli, 2014; Wilhelm et al., 2014). The proportion of release competent vesicles increases upon inactivity, contributing to the upscaling of presynaptic strengths (Thiagarajan et al., 2005; Menegon et al., 2006; Lazarevic et al., 2011; Verstegen et al., 2014). While *Bsn* is known to control the number of release-competent vesicles (Montenegro-Venegas et al., 2022), it has not yet been studied whether it is required for the inactivity-induced increase in synaptic release competence. To close this gap, we utilized the well-established synaptotagmin 1 (SYT1) antibody (SYT1Ab) uptake assay, which allows for quantitative assessment of SV fusion and retrieval at levels of individual synapses in cultured neurons. In this assay, the SYT1Ab recognizing the luminal domain of SV protein SYT1 was added to the culture media of living neurons to label exocytosed SV before retrieval by compensatory endocytosis (Kraszewski et al., 1995; Lazarevic et al., 2011; Guhathakurta et al., 2022). We analyzed SYT1Ab uptake in glutamatergic synapses, labeled for VGLUT1, in primary hippocampal neurons at basal conditions and upon a brief pulse application of 50mM KCl leading to depolarization-induced release of all release-capable SVs (total releasable pool, TRP) (Figure 4 A, C; Table 4), (Harata et al., 2001; Guhathakurta et al., 2022). The SYT1Ab uptake at basal and evoked (KCl-induced) conditions was increased by about 50% upon silencing in control cells (Figure 4B, D). In line with the previously reported effect of *Bsn* deletion on SV fusion competence in glutamatergic synapses, the SYT1Ab uptake was decreased by 45% at basal and by 33% at evoked conditions in Bsn^GT^ neurons as compared to Bsn^WT^ (Figure 4B, D) (Montenegro-Venegas et al., 2022). Importantly, silencing had no effect on SYT1Ab uptake in cells from *Bsn^GT^* mice revealing lack of inactivity-induced upscaling in release competence upon deletion of *Bsn* (Figure 4B, D).

**Figure 4:**
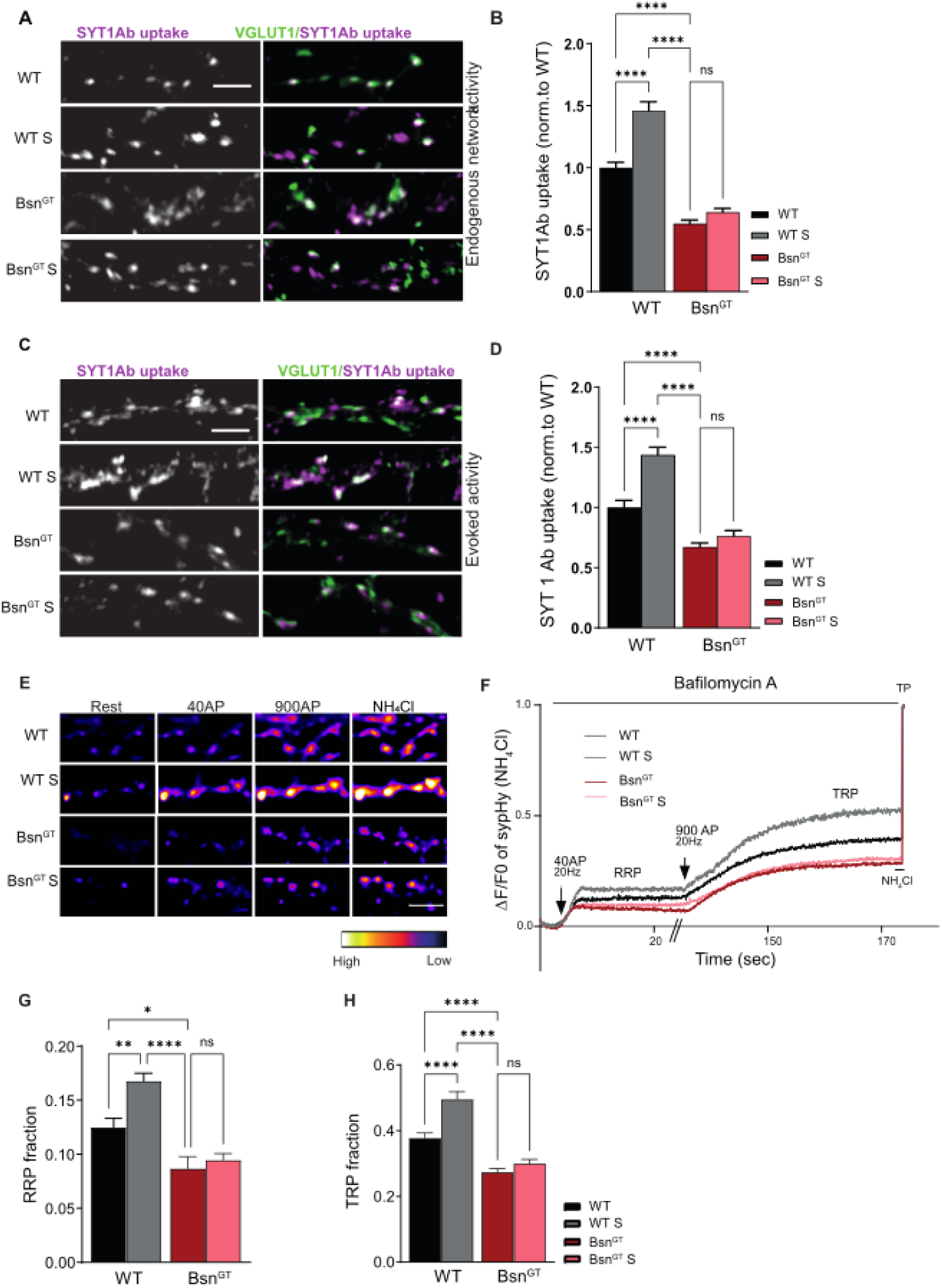
*Bsn* deletion impairs inactivity-induced homeostatic regulation of SV pools. (A, D) Representative images of SYT1Ab uptake and VGLUT1 immunoreactivity (to visualize excitatory boutons) from Bsn^WT^ and Bsn^GT^ (A, D) hippocampal neurons from control and silenced (S, treated with APV/CNQX) 20-21 DIV cells. Quantification of the SYT1Ab uptake at individual excitatory synapses during endogenous network activity conditions (A, B) or during depolarization with 50mM KCl for 4 minutes (C, D). Scale bar 3 μm. In plots, bars represent mean values obtained from 4 independent experiments. Whiskers, SEM is stated for each quantification. Data are shown as intensity values ± SEM normalized to the mean intensity value in the control group. (E) Representative images of Bsn^WT^ and Bsn^GT^ axons expressing sypHy at rest (left), at the peak of the response elicited by stimulation of 40APs (20Hz), 900 APs (20Hz) (middle), and after application of NH_4_Cl (right) that dequenches the sypHy probe and allows assessment of all SVs within synaptic termini. Experiment was performed in the presence of the proton pump inhibitor, bafilomycin A. Scale bar 2 µm (F). Representative average of traces of sypHyfluorescence plotted for Bsn^WT^ and Bsn^GT^ neurons treated with AP5/CNQX (gray and pink) (G, H). Data mean ± SEM of RRP and TRP fraction, respectively, calculated from the individual plateau values. Data are from 3 independent experiments. The statistic was assessed using two-way ANOVA with Tukey’s multiple comparison test; *p≤0.05, **p<0.01, ***p<0.001, ****p<0.0001, ns – not significant.

**Supplementary figure 4:**
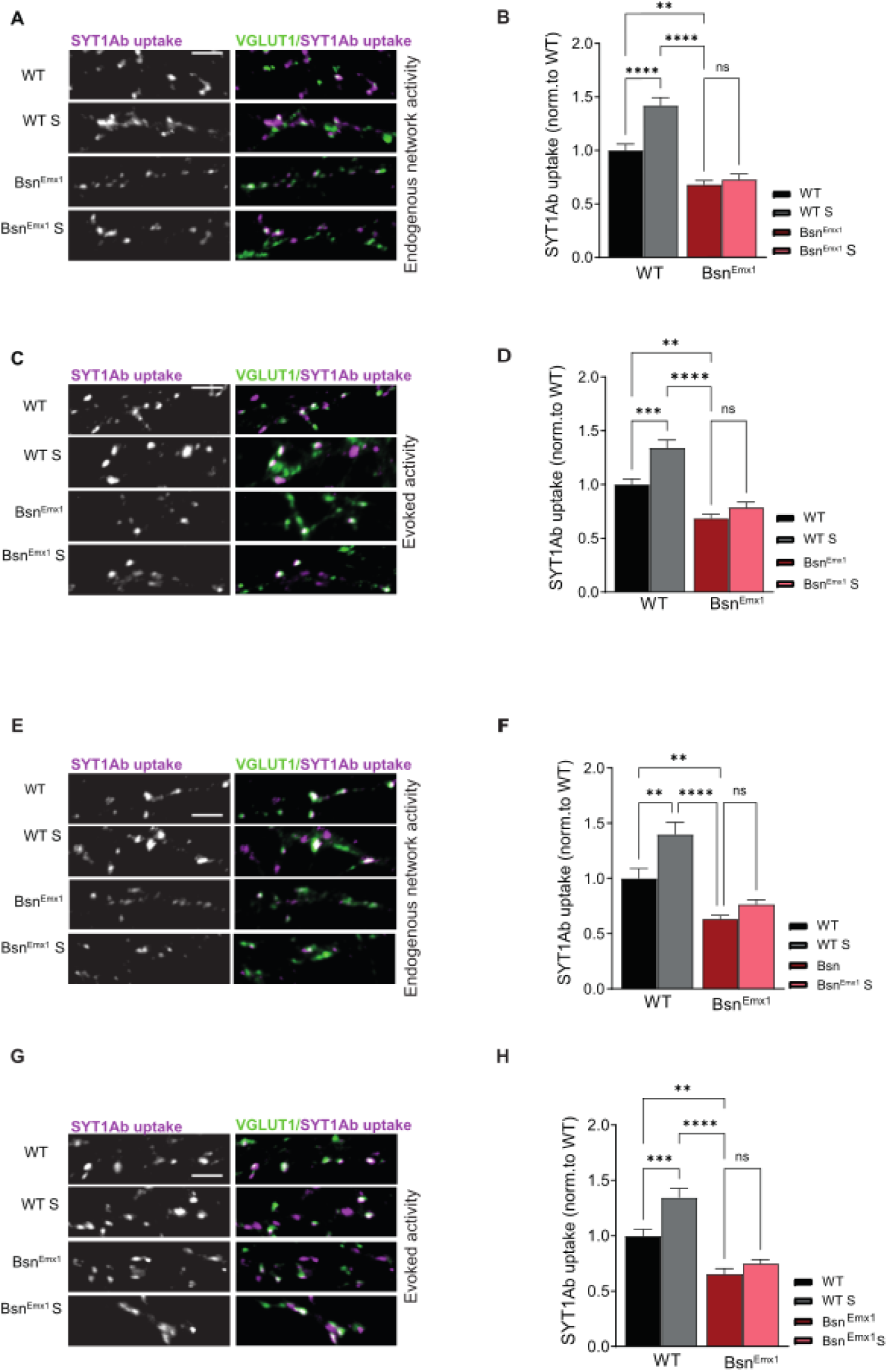
Defect in silencing-induced modulation of SV recycling in hippocampal and cortical neurons from Bsn^Emx1^ mice. (A, C, E, G) Representative images of SYT1Ab uptake and VGLUT1 immunoreactivity (to visualize excitatory boutons) from 20-21 DIV hippocampal (A, C) and cortical (E, G) neurons prepared from Bsn^WT^ or Bsn^Emx1^ animals and kept at basal conditions or with silencing (S) of network activity for 48 h. (B, D, F, H) Quantification of the SYT1Ab uptake at individual excitatory synapses driven by endogenous network activity (B, F) or upon depolarization with 50mM KCl for 4 minutes (D, H). Scale bar 3 μm. Data are shown as intensity values ± SEM normalized to the mean intensity value in the control group. In plots, bars represent mean values obtained from 3 independent experiments. The statistic was assessed using two-way ANOVA with Tukey’s multiple comparison test; *p≤0.05, **p<0.01, ***p<0.001, ****p<0.0001, ns – not significant.

To confirm the effect of silencing on synaptic vesicle pools by an independent method we imaged electrically evoked exocytosis of SV in Bsn^WT^ and Bsn^GT^ neurons using fluorescent synaptophysin-phluorin-tdimer2 (sypHy) reporter. This probe localizes to the membrane of SVs, with its pH-sensing domain in the lumen of SVs and tdimer2 red facing the cytoplasm, allowing us to identify the sypHy-expressing cells at rest. The fluorescence of sypHy is quenched in the acidic pH of SVs, emits fluorescence upon fusion of SVs with the plasma membrane, and is quenched again upon reacidification of retrieved SVs (Rose et al., 2013). The release of a RRP of SVs was triggered by the delivery of 40 pulses at 20 Hz, while all release-capable vesicles were released upon application of 900 pulses at 20 Hz (Burrone et al., 2006). In line with previously published data, we observed significantly decreased RRP and RP in untreated Bsn^GT^ neurons compared to Bsn^WT^ (Montenegro-Venegas et al., 2022). The 48h-long silencing increased RRP and TRP by about 30% in Bsn^WT^ but did not affect the size of RRP and TRP in neurons from Bsn^GT^ mice. This confirms defects in silencing-induced regulation of SV pool size in Bsn^GT^ animals (Figure 4E-H; Table 4).

Next, we tested whether the observations made in hippocampal neurons from constitutive Bsn^GT^ model could be replicated also in cultures from Bsn^Emx1^ mice with specific deletion of *Bsn* only in excitatory forebrain cells, in which we observed a defect in OD-plasticity *in vivo*. We prepared hippocampal neurons from Bsn^Emx1^ mice and induced silencing as described above. As in Bsn^GT^ neurons, silencing induced an increase in SYT1Ab uptake by about 42% at basal conditions and by about 36% upon 50mM KCl stimulation in control neurons, but no effect was evident in hippocampal neurons from Bsn^Emx1^ animals (Supplementary figure 4 A-D, Table S4). Finally, the same experiment was repeated using neurons prepared from cortices with the analogous outcome, confirming the universality of our finding for hippocampal and cortical neuronal networks *in vitro* (Supplementary figure 4E-H). Taken together, these experiments revealed that expression of *Bsn* in excitatory forebrain neurons is required for inactivity-induced regulation of SV pools.

### MD-induced changes in phosphorylation of SYN in V1 is disrupted in Bsn^Emx1^ mice

Changes in the phosphorylation of the synaptic protein SYN are a major molecular determinant of activity-dependent regulation of SV pool size (Cesca et al., 2010). Previous studies reveled aberrant phosphorylation of SYN upon *Bsn* deletion, which is caused by dysregulation of presynaptic CDK5/calcineurin and cAMP-PKA-PDE4 signaling axes and result in reduction of RRP and TRP pools (Montenegro-Venegas et al., 2022).

Changes in the phosphorylation status of SYN were reported for V1 of adult rats upon 3 day-long MD (Scott et al., 2010). To confirm that similar regulation also occurs in V1 of adult mice and to establish a time point suitable for investigating the phosphorylation status of SYN phospho-species in Bsn^Emx1^ animals, we first analyzed tissue lysates from contralateral and ipsilateral V1 of adult mice that were subjected to MD for either 3 or 7 days. Quantitative immunoblotting was performed using previously characterized phospho-specific antibodies against SYN phospho-species pS9SYN, pS62SYN, pS549SYN, pS553SYN and pS605SYN that detect these sites in SYN isoforms SYN1a, SYN1b, SYN2a, SYN3a and SYN2b (Supplementary figure 5A, B). Quantification revealed an increase in the relative abundance of phosphorylated SYN isoforms in V1 contralateral to the deprived eye, compared to the ipsilateral hemisphere. More pronounced effects were seen in animals that were subjected to MD for 7 days compared to animals analyzed only 3 days after MD (Supplementary figure 5C, D; Supplementary Table S5).

**Supplementary figure 5:**
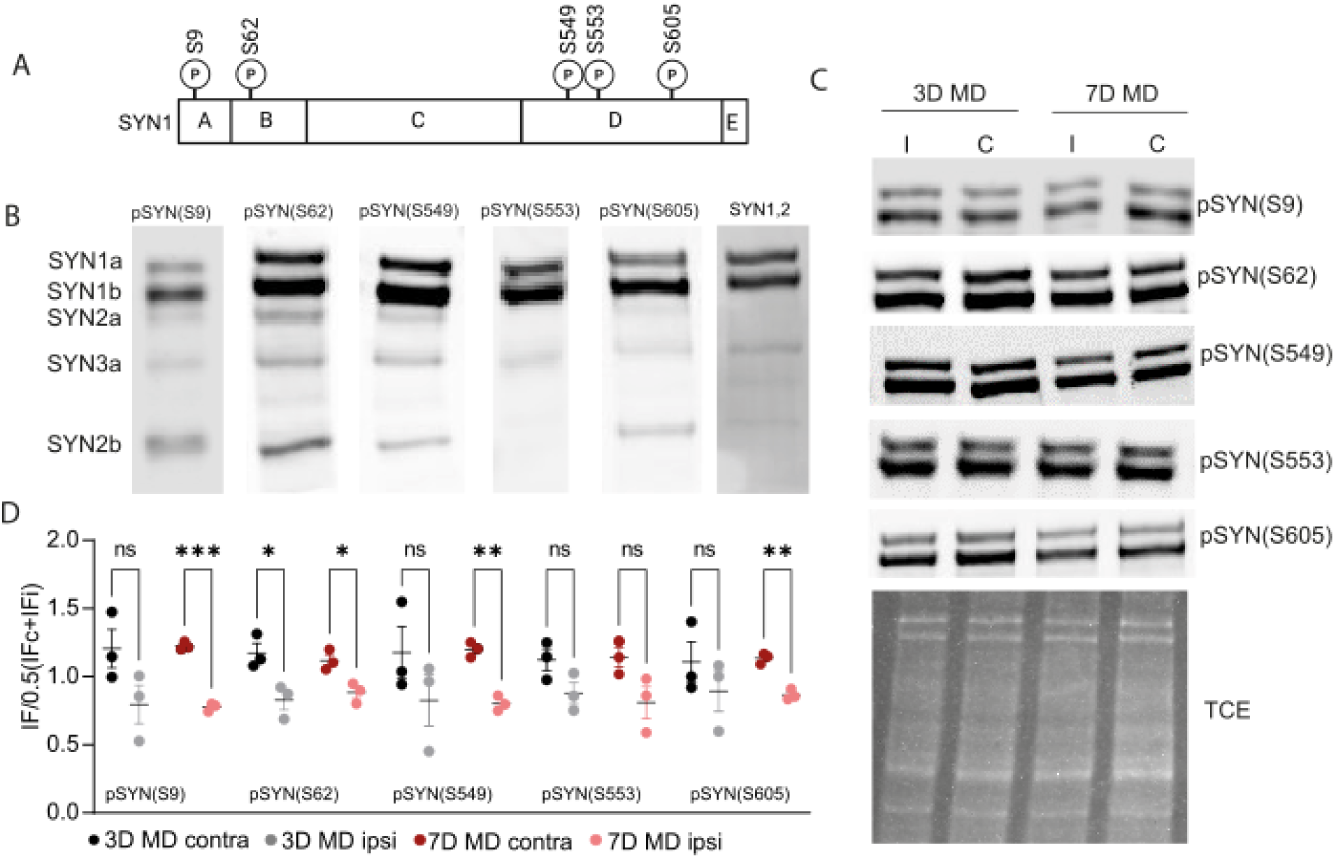
MD-induces dynamic changes in SYN phosphorylation in V1. **(A)** Schematic representation of the phosphorylation sites identified in the mammalian SYN isoforms. **(B)** Representative immunoblots depicting phosphorylation of detected SYN isoforms using the tested antibodies against pS9SYN, pS62SYN, pS549SYN, pS553SYN and pS605SYN phospho-species. **(C)** Representative immunoblots of contralateral (C/contra) and ipsilateral (I/ipsi) V1 tissue lysates from mice that underwent 3 days (3D; n=3) and 7 days (7D; n=3) of MD showing SYNI phosphorylation status for all SYN phospho-species tested. **(D)** Quantification of immunoblots shown in **(C)**. Data is expressed as mean ± SEM and significance were tested using Multiple unpaired t-test. Statistical significance is marked as *p < 0.05, **p < 0.01, and ***p < 0.001.

To test whether *Bsn* deletion affects these MD-induced changes in the phosphorylation of SYN, we analyzed samples from contralateral and ipsilateral V1 regions from Bsn^WT^ and Bsn^Emx1^ mice subjected to MD for 7 days. Animals with sham surgery and parallel handling were used as controls. We did not observe significant changes in the abundance of most SYN phospho-species between ipsilateral and contralateral V1 in sham treated animals of both genotypes (Figure 5A, B; Table 5). The quantifications revealed an increased relative abundance of the vast majority of the detected phospho-species of SYN isoforms in the lysates from contralateral V1 compared to ipsilateral V1 in Bsn^WT^ animals subjected 7d MD (Figure 5A, B; Table 5). The detected changes were highly significant for most detected phospho-species and SYN isoforms and reached relative changes by 20-65% (Table 5). In contrast, Bsn^Emx1^ animals subjected to 7d MD showed no difference in the phosphorylation status for any tested phospho-species of SYN isoforms and in fact had an overall tendency of phosphorylation being oppositely regulated between the V1 hemispheres compared to Bsn^WT^-mice (Figure 5A, B; Table 5). These data reveal that MD induces robust lateralized changes in SYN phosphorylation in V1. Moreover, they indicate that the MD-induced dynamic regulation of SYN phosphorylation states between the V1 hemispheres is lost in Bsn^Emx1^mice upon deletion of *Bsn* in neocortical glutamatergic neurons, that also failed to develop OD-plasticity upon MD.

**Figure 5:**
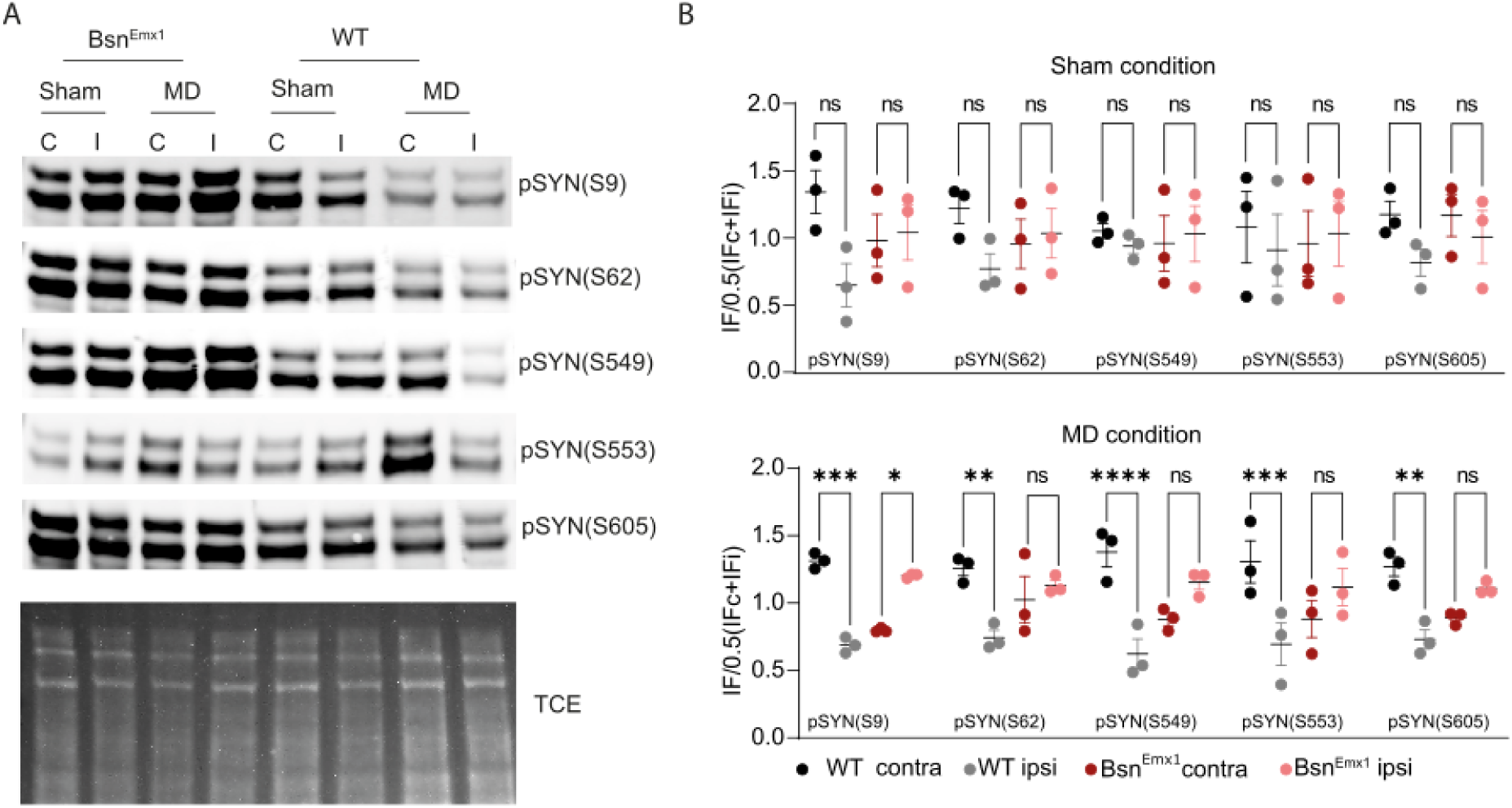
MD-induced changes in synapsin phosphorylation in V1 is abolished upon deletion of *Bsn* in neocortical glutamatergic neurons. **(A)** Representative immunoblots of contralateral (C/contra) and ipsilateral (I/ipsi) V1 tissue lysates from BsnEmx1 (n=3) and BsnWT (n=3) mice showing SYNI phosphorylation status for all SYN phospho-species tested, during sham and 7 days MD conditions respectively. **(B)** Quantification of immunoblots shown in **(A)**. Data is expressed as mean ± SEM and significance were tested using Multiple unpaired t-test with Holm-Sidak post-test. Statistical significance is marked as *p < 0.05, **p < 0.01, and ***p < 0.001.

## Discussion

In the present study, we analyzed the role of the presynaptic scaffolding protein BSN for experience-dependent changes in the mouse visual system. To this end, we used mouse lines with a global (constitutive) deletion of functional *Bsn* (Bsn^ΔEx4-5^ and Bsn^GT^ mice) and conditional strain with a *Bsn* deletion restricted to telencephalic excitatory neurons (Bsn^Emx1^ mice), and compared results to their respective Bsn^WT^ littermates. In Bsn^ΔEx4-5^ mice, we could confirm and extend the previously reported impairment in spatial vision in juvenile and adult Bsn^ΔEx4-5^ mice using the optomotor task (Goetze et al., 2010): both the baseline spatial frequency threshold (SFT) of the optomotor reflex and the experience-enabled improvement of reflex thresholds were severely compromised in animals with loss of functional *Bsn*. Furthermore, intrinsic signal optical imaging revealed that i) baseline ODIs were lower than in Bsn^WT^ control mice in both age groups and ii) OD-plasticity after brief monocular deprivation was present in juvenile but absent in adult animals. These observations suggested that that presynaptic protein BSN is essential for the expression of OD-plasticity in V1 of adult mice. This finding is striking since it points to a role of presynaptic mechanisms, which are underexplored in this process. The age-specific effect of *Bsn* mutation is in line with previous reports that showed differing mechanisms underlying adult and juvenile OD-plasticity in mouse V1 (Sato and Stryker, 2008; Ranson et al., 2012). However, constitutive deletion of *Bsn* induces progressive age-dependent retinal degeneration (Specht et al., 2007; Ryl et al., 2021) and it also leads to severe epileptiform seizures, which increase in severity and frequency as animals grow up (Altrock et al., 2003; Blondiaux et al., 2023). Seizures can cause extensive neuronal loss and severe brain damage (Olney et al., 1983; Sloviter, 1983, 1996). Thus, the physiological and morphological impairments due to the seizures might be more severe in adult than in juvenile mutant animals, which would fit to our data. Therefore, the absent OD-plasticity in adult Bsn^ΔEx4-5^ might have been caused by the increased epileptiform activity and not primarily by the deletion of functional *Bsn*. To decide between these alternatives, we additionally performed experiments in Bsn^Emx1^ mice with a deletion of *Bsn* restricted to telencephalic excitatory cells that show normal life expectancy and are not severely affected by epilepsy (Annamneedi et al., 2018; Blondiaux et al., 2023). Our additional experiments confirmed absent OD-plasticity in adult Bsn^Emx1^ mice further supporting our conclusion that *Bsn* is particularly important for adult OD-plasticity. We can also exclude age-dependent retinal degeneration and progressive loss of vision observed in Bsn^ΔEx4-5^ as an underlying cause of the compromised adult OD-plasticity since our optomotor measurements clearly showed normal basic spatial vision and normal experience-enabled vision enhancements in Bsn^Emx1^ mice. In fact, we had expected normal optomotor results that primarily depend on function of retinal or subcortical neurons in the Bsn^Emx1^ mice, as *Emx1* promoter is not active in these regions. In contrast, confirming previous data (Goetze et al., 2010), basic spatial vision was severely compromised in both juvenile and adult Bsn^ΔEx4-5^ mice. In addition, experience-enabled vision enhancements were also reduced in Bsn^ΔEx4-5^ mice compared to their Bsn^WT^ littermates. We hypothesize that these impairments are due to the defects in the subcortical and retinal circuitry caused by the constitutive lack of functional *Bsn* independently of age. Thus, our data thus clearly show that presynaptic protein BSN is essential for the expression of OD-plasticity in V1 of adult mice.

The MD-induced OD-plasticity differs qualitatively and mechanistically between juvenile and adult mice (Desai et al., 2002; Heynen et al., 2003; Lehmann and Löwel, 2008; Sato and Stryker, 2008; Ranson et al., 2012; Stryker and Löwel, 2018). While in juvenile animals, 4 days of pattern vision deprivation (MD) causes a reduction in V1-activation mediated by visual stimulation of the closed in adults, OD-shifts after MD are mainly driven by strengthening of V1-activation via stimulating the open eye (Desai et al., 2002; Lehmann and Löwel, 2008; Sato and Stryker, 2008; Espinosa and Stryker, 2012). Our experiments revealed that deletion of *Bsn* did not interfere with juvenile OD-plasticity, i.e. the depression of V1-activation after visual stimulation of the previously deprived (contralateral) eye. In contrast, deletion of *Bsn* from telencephalic glutamatergic neurons completely abolished the MD-induced strengthening of V1-activation via the open eye in adults. Considering the selective presynaptic localization of BSN and the fact that its deletion in neocortical excitatory neurons was sufficient to abolish adult MD-induced OD plasticity, we propose the existence of BSN-dependent presynaptic plasticity mechanisms in neocortical glutamatergic neurons, which critically contributes to the MD-induced OD plasticity in adults.

Both homeostatic and Hebbian synaptic plasticity mechanisms cooperate during OD-plasticity in V1. While LTP was largely unaffected in Bsn^Emx1^ animals (Annamneedi et al., 2018) the effect of *Bsn* deletion on presynaptic homeostatic plasticity was unclear and therefore addressed in this study. Using patch clamp recordings and immunostainings in cultured primary hippocampal and cortical neurons with *Bsn* deletion, we confirmed a normal silencing-induced increase in the amplitude of mEPSCs mediated by an enhanced externalization of AMPA receptors in dendritic spines. In contrast, the silencing of network activity failed to increase the frequency of mEPSCs, which is mediated by an increased rate of neurotransmitter release from the presynapse upon *Bsn* deletion, confirming a specific defect in *presynaptic* homeostatic scaling. Subsequent live-imaging experiments revealed that the size of readily releasable and recycling pools of SVs, which is normally increased upon silencing, did not adapt to the network activity and remained significantly reduced upon deletion of *Bsn*. Previous studies showed that aberrant activity of presynaptic kinases and phosphatases causes defects in SV release in the absence of *Bsn*, with aberrant phosphorylation of SYN and changes in size of SV pools and SV release competence (Montenegro-Venegas et al., 2022). Interestingly in this context, MD-induced changes of SYN phosphorylation in V1 have been documented (Scott et al., 2010; Fu et al., 2015). Scott and coauthors reported an increased phosphorylation of site 1 (S9) and site 3 (S604) of SYN in samples of binocular V1 ipsilateral to the deprived eye 3 days after MD in adult rats (P70-90), while no changes were evident in juveniles (P28) (Scott et al., 2010). In contrast, Fu and coauthors, described increased site 1 phosphorylation 7 days after MD in contralateral V1 of juvenile (P28 and P35), but not young adult (P42 or P60) mice (Fu et al., 2015). We observed increased SYN phosphorylation in V1 contralateral to the deprived eye in adult animals 3 and 7 days after MD. These differences may be due to the different species, timing or experimental protocols, nevertheless from all these studies it is evident that MD affects SYN phosphorylation in V1. SYN plays a key role in SV clustering within presynaptic boutons. SYN phosphorylation controls the mobility of SVs and their availability for evoked fusion and is therefore a key regulator of efficient neurotransmitter release and is under tight control during various types of presynaptic plasticity (Cesca et al., 2010). Thus, changes in SYN phosphorylation status directly indicate adaptation of presynaptic properties to changed input upon MD. Strikingly, the MD-induced shift in the SYN phosphorylation was completely abolished in the Bsn^Emx1^ mice that also failed to show any MD-induced OD-shift. Confirming previous work (Montenegro-Venegas et al., 2022), our data (Figure 5A) also showed changes in SYN phosphorylation in cortical samples prepared from naïve mice with *Bsn* deletion which is in line with a pivotal role of *Bsn* in the regulation of dynamic SYN phosphorylation *in vivo*. Based on all these observations, we propose that *Bsn* deletion disrupts MD-induced OD-plasticity because it interferes with MD-induced changes in SYN phosphorylation, which are known to control SV dynamics and thereby presynaptic strengths (Cesca et al., 2010)

Additional mechanisms not specifically addressed in this study might also contribute to the defects in adult MD-induced OD-plasticity in *Bsn* mutants. In Bsn^ΔEx4-5^ mice, progressively increasing brain-derived growth factor (BDNF) and neurotrophic tyrosine kinase, receptor, type 2 (TRKB) levels were detected in brain tissues, although it was not possible to delineate whether this increase causes or arise from seizure activity in these animals (Ghiglieri et al., 2010; Heyden et al., 2011; Dieni et al., 2012). While adult Bsn^Emx1^ mice do not show epileptic seizures, they exhibit enhanced hippocampal excitability due to increased BDNF/TRKB signaling, as demonstrated by successful reversal of this phenotype by acute inhibition of TRKB receptor (Annamneedi et al., 2018; Annamneedi et al., 2021). Enhanced BDNF/TRKB signaling promotes synaptic structural and functional plasticity and enables OD-plasticity in older (>P100) adult animals (Winkel et al., 2021). BDNF/TRKB signaling was also shown to promote maturation of inhibitory cells in neocortex, which coincides with the termination of the critical period for OD-plasticity (Huang et al., 1999; Kaneko et al., 2012). On the other hand, activation of TrkB receptors on parvalbumin positive cells was necessary for reinstatement of OD-plasticity in adult animals upon treatment with antidepressant or digestion of extracellular matrix indicating a more complex role of TrkB signaling in regulating inhibitory drive (Winkel et al., 2021). Interestingly, TRKB signaling was also shown to activate RAS/MAPK signaling to increase SYN phosphorylation (Jovanovic et al., 1996) as observed in Bsn^Emx1^ mice in our study. Therefore, it is possible that increased BDNF/TRKB activity could contribute to the absent OD-plasticity of adult Bsn^Emx1^ animals, however deciphering the exact cellular mechanisms and pathways will need further research.

## Supporting information

Supplemetary tables: datasets, statistics

## Author contributions

Conceptualization: A.F., S.L., E.D.G., C.M.-V., D.G, B.G., F.G.

Methodology: C.M.-V., D.G., B.G. F.G., M.H., S.L., A.F.,

Investigation: C.M.-V., D.G., B.G., J.B., F.G., S.P., E.P.-F.,

Formal analysis: C.S., C.M.-V., D.G., S.P., E.P.-F., A.F.,

Validation: C.S., C.M.-V., D.G., A.F.,

Visualization: C.S., C.M.-V., D.G., B.G., J.B., A.F., S.L.

Writing—original draft: A.F., C.S., C.M.-V., D.G., B.G., S.L.

Writing—review and editing: all authors

Resources: M.H., E.D.G., S.L., A.F.

Funding acquisition: A.F., E.D.G., C.M.-V., A.A., S.L.,

Project administration: A.F., S.L.

Supervision: M.H., A.F., E.D.G., S.L.

## Acknowledgment

We thank Juliana Monti, Kati Ebert, and the team of the PETZ and LIN animal facility for excellent technical support and Stefanie Kempf, Silvia Bose, Kalina Makowiecki and Merle Fricke for support during data acquisition and analyses.

## Funding

The research was funded by DFG FE1335/3, DFG MH3604/11, CRC 779 to E.D.G. and A.F., and CRC889, Project B5 to S.L.

Center for Behavioral Brain Sciences—CBBS promoted by Europäische Fonds für regionale Entwicklung—EFRE (ZS/2016/04/78113) to CMV and AA.

The Leibniz Graduate School “SynaptoGenetics” (Leibniz SAW program) to E.D.G and A.F.

