## Supplementary material for "Bassoon is required for adult mouse ocular dominance plasticity and experience-induced changes in synapsin phosphorylation in visual cortex": Supplemetary tables: datasets, statistics

**Table 2**

### ODI

| Figure | genotype | treatment | Mean ± SEM | Samples | 2-way ANOVA results | F(df) | p-value |
|---|---|---|---|---|---|---|---|
| **Figure 2B** | **BsnWT** | noMD | 0.19 ± 0.01 | 7 | genotype | 7.0 (1, 30) | **0,013** |
| | | MD | 0.08 ± 0.03 | 8 | treatment | 4.3 (1, 30) | **0,048** |
| | **BsnEmx1** | noMD | 0.21 ± 0.02 | 7 | treatment x genotype | 3.6 (1, 30) | n.s. |
| | | MD | 0.20 ± 0.03 | 12 | | | |

| FDR corrected posthoc comparisson | p-value |
|---|---|
| noMD:Adult BsnWT vs. noMD:Adult BsnEmx1 | 0,8 |
| noMD:Adult BsnWT vs. 7d MD:Adult BsnWT | **0,0233** |
| noMD:Adult BsnWT vs. 7d MD:Adult BsnEmx1 | 0,8 |
| noMD:Adult BsnEmx1 vs. 7d MD:Adult BsnWT | **0,0097** |
| noMD:Adult BsnEmx1 vs. 7d MD:Adult BsnEmx1 | 0,903 |
| 7d MD:Adult BsnWT vs. 7d MD:Adult BsnEmx1 | **0,0089** |

### V1-activity

| Figure | genotype | treatment / eye | Mean ± SEM | Samples | 3-way ANOVA results | F(df) | p-value |
|---|---|---|---|---|---|---|---|
| **Figure 2B** | **BsnWT** | noMD / contra | 1.2 ± 0.1 | 7 | eye | 60.93 (1, 30) | **<0.0001** |
| | | noMD / ipsi | 0.9 ± 0.1 | 7 | genotype | 0.2709 (1, 30) | n.s. |
| | | MD / contra | 1.2 ± 0.1 | 8 | treatment | 0.1707 (1, 30) | n.s. |
| | | MD / ipsi | 1.0 ± 0.1 | 8 | eye x genotype | 2.890 (1, 30) | n.s. |
| | **BsnEmx1** | noMD / contra | 1.3 ± 0.2 | 7 | eye x treatment | 0.6494 (1, 30) | n.s. |
| | | noMD / ipsi | 0.9 ± 0.1 | 7 | genotype x treatment | 0.1 (1, 30) | n.s. |
| | | MD / contra | 1.3 ± 0.1 | 12 | eye x genotype x treatment | 0.7 (1, 30) | n.s. |
| | | MD / ipsi | 0.9 ± 0.1 | 12 | | | |

| FDR corrected posthoc comparisson | p-value |
|---|---|
| Contra:BsnWT noMD vs. Ipsi:BsnWT noMD | **0,0044** |
| Contra:BsnWT MD vs. Ipsi:BsnWT MD | 0,1046 |
| Contra:BsnEmx1 noMD vs. Ipsi:BsnEmx1 noMD | **0,0008** |
| Contra:BsnEmx1 MD vs. Ipsi:BsnEmx1 MD | **<0.0001** |
| Contra:BsnWT noMD vs. Contra:BsnWT MD | 0,9422 |
| Contra:BsnEmx1 noMD vs. Contra:BsnEmx1 MD | 0,9422 |
| Ipsi:BsnWT noMD vs. Ipsi:BsnWT MD | 0,8393 |
| Ipsi:BsnEmx1 noMD vs. Ipsi:BsnEmx1 MD | 0,9422 |
| Contra:BsnWT noMD vs. Contra:BsnEmx1 noMD | 0,8575 |
| Contra:BsnWT MD vs. Contra:BsnEmx1 MD | 0,8393 |
| Ipsi:BsnWT noMD vs. Ipsi:BsnEmx1 noMD | 0,9422 |
| Ipsi:BsnWT MD vs. Ipsi:BsnEmx1 MD | 0,9422 |

### Spatial frequency threshold (SFT)

| Figure | genotype | treatment / MD-day | Mean ± SEM | Samples | 3-way ANOVA | F(df) | p-value |
|---|---|---|---|---|---|---|---|
| **Figure 2C/E** | **BsnWT** | noMD / D0 | 0.40 ± 0.00 | 6 | genotype | 0.2 (1,22) | n.s. |
| | | noMD / D7 | 0.42 ± 0.01 | 6 | treatment | 28.2 (1,22) | **<0.0001** |
| | | MD / D0 | 0.40 ± 0.01 | 7 | MD-day | 116.9 (1,22) | **<0.0001** |
| | | MD / D7 | 0.49 ± 0.00 | 7 | treatment x MD-day | 50.0 (1,22) | **<0.0001** |
| | **BsnEmx1** | noMD / D0 | 0.39 ± 0.02 | 5 | MD-day x genotype | 4.7 (1,22) | **0,041** |
| | | noMD / D7 | 0.42± 0.01 | 5 | treatment x genotype | 0.4 (1,22) | n.s. |
| | | MD / D0 | 0.39 ± 0.01 | 8 | MD-day x genotype x treatment | 1.5 (1,22) | n.s. |
| | | MD / D7 | 0.51 ± 0.01 | 8 | | | |

| FDR corrected posthoc comparisson | p-value |
|---|---|
| D0:BsnWT noMD vs. D7:BsnWT noMD | 0,92 |
| D0:BsnWT 7dMD vs. D7:BsnWT 7dMD | **<0.0001** |
| D0:BsnEmx1 noMD vs. D7:BsnEmx1 noMD | 0,4446 |
| D0:BsnEmx1 7dMD vs. D7:BsnEmx1 7dMD | **<0.0001** |
| D0:BsnWT noMD vs. D0:BsnWT 7dMD | >0.9999 |
| D0:BsnEmx1 noMD vs. D0:BsnEmx1 7dMD | >0.9999 |
| D7:BsnWT noMD vs. D7:BsnWT 7dMD | **<0.0001** |
| D7:BsnEmx1 noMD vs. D7:BsnEmx1 7dMD | **<0.0001** |
| D0:BsnWT noMD vs. D0:BsnEmx1 noMD | 0,9999 |
| D0:BsnWT 7dMD vs. D0:BsnEmx1 7dMD | 0,9842 |
| D7:BsnWT noMD vs. D7:BsnEmx1 noMD | >0.9999 |
| D7:BsnWT 7dMD vs. D7:BsnEmx1 7dMD | 0,4448 |

### gain on baseline spatial frequency threshold (SFT)

| Figure | genotype | treatment / MD-day | Mean ± SEM | Samples | 3-way ANOVA | F(df) | p-value |
|---|---|---|---|---|---|---|---|
| **Figure 2D** | **BsnWT** | noMD / D7 | 3.7 ± 1.4 | 6 | genotype | 0.6 (1, 22) | ns |
| | | MD / D7 | 11.4 ± 0.8 | 7 | treatment | 43.2 (1, 22) | **<0.0001** |
| | **BsnEmx1** | noMD / D7 | 2.8 ± 1.1 | 5 | MD-day | 116 (2.3, 50.5) | **<0.0001** |
| | | MD / D7 | 12.7 ± 0.9 | 8 | treatment x MD-day | 40.8 (7, 154) | **<0.0001** |
| | | | | | MD-day x genotype | 0.78 (7, 154) | n.s. |
| | | | | | treatment x genotype | 0.4 (7, 154) | n.s. |
| | | | | | MD-day x genotype x treatment | 1.1 (1, 154) | n.s. |

**Table 1**

#### ODI

| Figure | genotype | treatment | SEM | Samples | 2-way ANOVA results | F(df) | p-value |
|---|---|---|---|---|---|---|---|
| **Figure 1C** | **BsnWT** | noMD | 0.25 ± 0.03 | 5 | genotype | 0.2 (1, 17) | n.s. |
| P25-35 | | MD | -0.05 ± 0.03 | 3 | treatment | 56.6 (1, 17) | **<0.0001** |
| | **BsnΔEx4-5** | noMD | 0.15 ± 0.02 | 8 | treatment x genotype | 8.2 (1, 17) | **0,011** |
| | | MD | 0.02 ± 0.03 | 5 | | | |
| **Figure 1D** | **BsnWT** | noMD | 0.23 ± 0.03 | 11 | genotype | 0.8 (1, 34) | n.s. |
| P50-95 | | MD | -0.03 ± 0.04 | 8 | treatment | 15.8 (1, 34) | **0,0003** |
| | **BsnΔEx4-5** | noMD | 0.12 ± 0.02 | 11 | treatment x genotype | 20.5 (1, 34) | **<0.0001** |
| | | MD | 0.14 ± 0.03 | 6 | | | |

| FDR corrected posthoc comparisson | p-value |
|---|---|
| noMD:Juv. BsnWT vs. noMD:Juv. BsnΔEx4-5 | **0,017** |
| noMD:Juv. BsnWT vs. 7d MD:Juv. BsnWT | **<0.0001** |
| noMD:Juv. BsnWT vs. 7d MD:Juv. BsnΔEx4-5 | **<0.0001** |
| noMD:Juv. BsnΔEx4-5 vs. 7d MD:Juv. BsnWT | **0,0003** |
| noMD:Juv. BsnΔEx4-5 vs. 7d MD:Juv. BsnΔEx4-5 | **0,0022** |
| 7d MD:Juv. BsnWT vs. 7d MD:Juv. BsnΔEx4-5 | 0,1517 |
| noMD:Adult BsnWT vs. noMD:Adult BsnΔEx4-5 | **0,004** |
| noMD:Adult BsnWT vs. 7d MD:Adult BsnWT | **<0.0001** |
| noMD:Adult BsnWT vs. 7d MD:Adult BsnΔEx4-5 | 0,0548 |
| noMD:Adult BsnΔEx4-5 vs. 7d MD:Adult BsnWT | **0,0028** |
| noMD:Adult BsnΔEx4-5 vs. 7d MD:Adult BsnΔEx4-5 | 0,7154 |
| 7d MD:Adult BsnWT vs. 7d MD:Adult BsnΔEx4-5 | **0,004** |

#### V1-activity

| Figure | genotype | treatment / eye | SEM | Samples | 3-way ANOVA results | F(df) | p-value |
|---|---|---|---|---|---|---|---|
| **Figure 1C** | **BsnWT** | noMD / contra | 2.2 ± 0.4 | 5 | eye | 15.0 (1, 17) | **0,0012** |
| **P25-35** | | noMD / ipsi | 1.3 ± 0.2 | 5 | genotype | 0.1 (1, 17) | n.s. |
| | | MD / contra | 1.5 ± 0.1 | 3 | treatment | 0.8 (1, 17) | n.s. |
| | | MD / ipsi | 1.5 ± 0.1 | 3 | eye x genotype | 1.5 (1, 17) | n.s. |
| | **BsnΔEx4-5** | noMD / contra | 2.1 ± 0.3 | 8 | eye x treatment | 16.4 (1, 17) | **0,0008** |
| | | noMD / ipsi | 1.6 ± 0.3 | 8 | genotype x treatment | 0.01 (1, 17) | n.s. |
| | | MD / contra | 1.5 ± 0.1 | 5 | eye x genotype x treatment | 0.7 (1, 17) | n.s. |
| | | MD / ipsi | 1.6 ± 0.1 | 5 | | | |
| **Figure 1D** | **BsnWT** | noMD / contra | 2.2 ± 0.2 | 11 | eye | 25.26 (1, 32) | **<0.0001** |
| **P50-95** | | noMD / ipsi | 1.3 ± 0.1 | 11 | genotype | 0.7484 (1, 32) | n.s. |
| | | MD / contra | 2.6 ± 0.3 | 8 | treatment | 0.4443 (1, 32) | n.s. |
| | | MD / ipsi | 2.7 ± 0.3 | 8 | eye x genotype | 1.808 (1, 32) | n.s. |
| | **BsnΔEx4-5** | noMD / contra | 2.4 ± 0.4 | 11 | eye x treatment | 2.181 (1, 32) | n.s. |
| | | noMD / ipsi | 1.9 ± 0.4 | 11 | genotype x treatment | 3.170 (1, 32) | 0,085 |
| | | MD / contra | 2.2 ± 0.3 | 6 | eye x genotype x treatment | 10.05 (1, 32) | **0,003** |
| | | MD / ipsi | 1.4 ± 0.2 | 6 | | | |

| FDR corrected posthoc comparisson | p-value |
|---|---|
| Contra:juv BsnWT noMD vs. Ipsi:juv BsnWT noMD | **0,001** |
| Contra:juv BsnWT MD vs. Ipsi:juv BsnWT MD | 0,9949 |
| Contra:juv BsnΔEx4-5 noMD vs. Ipsi:juv BsnΔEx4-5 noMD | **0,0075** |
| Contra:juv BsnΔEx4-5 MD vs. Ipsi:juv BsnΔEx4-5 MD | 0,9949 |
| Contra:juv BsnWT noMD vs. Contra:juv BsnWT MD | 0,5345 |
| Contra:juv BsnΔEx4-5 noMD vs. Contra:juv BsnΔEx4-5 MD | 0,5345 |
| Ipsi:juv BsnWT noMD vs. Ipsi:juv BsnWT MD | 0,9949 |
| Ipsi:juv BsnΔEx4-5 noMD vs. Ipsi:juv BsnΔEx4-5 MD | 0,9949 |
| Contra:juv BsnWT noMD vs. Contra:juv BsnΔEx4-5 noMD | 0,9949 |
| Contra:juv BsnWT MD vs. Contra:juv BsnΔEx4-5 MD | 0,9949 |
| Ipsi:juv BsnWT noMD vs. Ipsi:juv BsnΔEx4-5 noMD | 0,9949 |
| Ipsi:juv BsnWT MD vs. Ipsi:juv BsnΔEx4-5 MD | 0,9949 |
| Contra:adult BsnWT noMD vs. Ipsi:adult BsnWT noMD | **0,0002** |
| Contra:adult BsnWT MD vs. Ipsi:adult BsnWT MD | 0,6872 |
| Contra:adult BsnΔEx4-5 noMD vs. Ipsi:adult BsnΔEx4-5 noMD | **0,0337** |
| Contra:adult BsnΔEx4-5 MD vs. Ipsi:adult BsnΔEx4-5 MD | **0,0103** |
| Contra:adult BsnWT noMD vs. Contra:adult BsnWT MD | 0,556 |
| Contra:adult BsnΔEx4-5 noMD vs. Contra:adult BsnΔEx4-5 MD | 0,6872 |
| Ipsi:adult BsnWT noMD vs. Ipsi:adult BsnWT MD | **0,0195** |
| Ipsi:adult BsnΔEx4-5 noMD vs. Ipsi:adult BsnΔEx4-5 MD | 0,4954 |
| Contra:adult BsnWT noMD vs. Contra:adult BsnΔEx4-5 noMD | 0,6872 |
| Contra:adult BsnWT MD vs. Contra:adult BsnΔEx4-5 MD | 0,5782 |
| Ipsi:adult BsnWT noMD vs. Ipsi:adult BsnΔEx4-5 noMD | 0,3067 |
| Ipsi:adult BsnWT MD vs. Ipsi:adult BsnΔEx4-5 MD | **0,0459** |

#### Spatial frequency threshold (SFT)

| Figure | genotype | treatment / MD-day | Mean ± SEM | Samples | repeated measures 2-way ANOVA | F(df) | p-value |
|---|---|---|---|---|---|---|---|
| **Figure 1E/G** | **BsnWT** | MD / D0 | 0.39 ± 0.00 | 6 | genotype | 2157 (1,17) | **<0.0001** |
| P25-35 | | MD / D4 | 0.46 ± 0.00 | 6 | MD-day | 491.9 (1,17) | **<0.0001** |
| | **BsnΔEx4-5** | MD / D0 | 0.20 ± 0.00 | 13 | MD-day x genotype | 162.1 (1,17) | **<0.0001** |
| | | MD / D4 | 0.22 ± 0.00 | 13 | | | |
| **Figure 1H/J** | **BsnWT** | MD / D0 | 0.40 ± 0.00 | 9 | genotype | 5849 (1,13) | **<0.0001** |
| P50-95 | | MD / D7 | 0.50 ± 0.00 | 9 | MD-day | 314.4 (1,13) | **<0.0001** |
| | **BsnΔEx4-5** | MD / D0 | 0.20 ± 0.00 | 6 | MD-day x genotype | 87.4 (1,13) | **<0.0001** |
| | | MD / D7 | 0.24 ± 0.00 | 6 | | | |

| FDR corrected posthoc comparisson | p-value |
|---|---|
| BsnWT D0 vs D4 | **<0.0001** |
| BsnΔEx4-5 D0 vs D4 | **<0.0001** |
| BsnWT D0 vs BsnΔEx4-5 D0 | **<0.0001** |
| BsnWT D4 vs BsnΔEx4-5 D4 | **<0.0001** |
| BsnWT D0 vs D7 | **<0.0001** |
| BsnΔEx4-5 D0 vs D7 | **0,0002** |
| BsnWT D0 vs BsnΔEx4-5 D0 | **<0.0001** |
| BsnWT D7 vs BsnΔEx4-5 D7 | **<0.0001** |

#### gain on baseline spatial frequency threshold (SFT)

| Figure | genotype | treatment / MD-day | Mean ± SEM | Samples | repeated measures 2-way ANOVA | F(df) | p-value |
|---|---|---|---|---|---|---|---|
| **Figure 1F** | **BsnWT** | MD / D4 | 18 ± 1 % | 6 | genotype | 8.1 (1,17) | **0,011** |
| P25-35 | **BsnΔEx4-5** | MD / D4 | 9 ± 1 % | 13 | MD-day | 135 (1.8, 29.8) | **<0.0001** |
| | | | | | MD-day x genotype | 13.3 (4,68) | **<0.0001** |
| **Figure 1I** | **BsnWT** | MD / D7 | 27 ± 1 % | 9 | genotype | 13.9 (1,13) | **0,003** |
| P50-95 | **BsnΔEx4-5** | MD / D7 | 16 ± 3 % | 6 | MD-day | 55.3 (2.7, 35.2) | **<0.0001** |
| | | | | | MD-day x genotype | 4.3 (7,91) | **<0.0001** |

**Table 3**

**Electrophysiology**

| Figure | genotype | treatment | Mean ± SEM | samples/ independent experiments | statistical test | F(df) | p-value |
|---|---|---|---|---|---|---|---|
| | | | | | **2-way ANOVA** | | |
| **Figure 3C** | **WT** | no treatment | 530 ± 68.96 | 10 / 3 | interaction | 3.065 (1, 38) | 0,0881 |
| Inter-event interval | | AP5/CNQX | 113 ± 18.72 | 12 / 3 | treatment | 5.424 (1, 38) | **0,0253** |
| | **BsnGT** | no treatment | 1187 ±108.2 | 10 / 3 | genotype | 66.99 (1, 38) | **<0.0001** |
| | | AP5/CNQX | 1128 ± 170.3 | 10 / 3 | | | |
| **Figure 3E** | **WT** | no treatment | 20.09 ± 1.48 | 10 / 3 | interaction | 0.5732 (1, 38) | 0,4537 |
| Amplitude | | AP5/CNQX | 32.64 ± 3.11 | 12 / 3 | treatment | 15.47 (1, 38) | **0,0003** |
| | **BsnGT** | no treatment | 20.13 ± 1.92 | 10 / 3 | genotype | 0.5490 (1, 38) | 0,463 |
| | | AP5/CNQX | 28.63 ± 3.40 | 10 / 3 | | | |
| **Surface pan-GluA** | | | | | **2-way ANOVA** | | |
| **Figure 3G** | **WT** | no treatment | 1.000 ± 0.015 | 38 / 3 | interaction | 0.738 (1, 20) | 0.4003 |
| | **WT** | AP5/CNQX | 1.162 ± 0.027 | 37 / 3 | treatment | 35.20 (1, 20) | **<0.0001** |
| | **BsnKO** | no treatment | 0.975 ± 0.021 | 37 / 3 | genotype | 3.568 (1, 20) | 0.073 |
| | **BsnKO** | AP5/CNQX | 1.097 ± 0.029 | 37 / 3 | | | |

**Table 4**

| Figure | genotype | treatment | Mean ± SEM | samples/ independent experiments | statistical test | F(df) | p-value |
|---|---|---|---|---|---|---|---|
| **Syt1 uptake assay** | | | | | **2-way ANOVA** | | |
| **Figure 4B** | **WT** | no treatment | 1.00 ± 0.04 | 39 / 4 | interaction | 12.25 (1, 188) | **0,0006** |
| Endogenous activity | | AP5/CNQX | 1.46 ± 0.07 | 57 / 4 | treatment | 27.83 (1, 188) | **<0.0001** |
| | **BsnGT** | no treatment | 0.55 ± 0.03 | 42 / 4 | genotype | 148.0 (1, 188) | **<0.0001** |
| | | AP5/CNQX | 0.64 ± 0.03 | 54 / 4 | | | |
| **Figure 4D** | **WT** | no treatment | 1.00 ± 0.06 | 42 / 4 | interaction | 11.36 (1, 180) | **0,0009** |
| Evoked activity | | AP5/CNQX | 1.44 ± 0.06 | 47 / 4 | treatment | 26.36 (1, 180) | **<0.0001** |
| | **BsnGT** | no treatment | 0.67 ± 0.03 | 47 / 4 | genotype | 95.09 (1, 180) | **<0.0001** |
| | | AP5/CNQX | 0.76 ± 0.05 | 48 / 4 | | | |
| **Vesicle Pool** | | | | | **2-way ANOVA** | | |
| **Figure 4G** | **WT** | no treatment | 0.13 ± 0.009 | 15 / 3 | interaction | 3.946 (1, 53) | 0,0522 |
| RRP | | AP5/CNQX | 0.17 ± 0.007 | 12 / 3 | treatment | 7.908 (1, 53) | **0,0069** |
| | **BsnGT** | no treatment | 0.087 ± 0.011 | 16 / 3 | genotype | 38.72 (1, 53) | **<0.0001** |
| | | AP5/CNQX | 0.095 ± 0.006 | 14 / 3 | | | |
| **Figure 4H** | **WT** | no treatment | 0.38 ± 0.016 | 15 / 3 | interaction | 8.607 (1, 53) | **0,0049** |
| TRP | | AP5/CNQX | 0.50 ± 0.023 | 12 / 3 | treatment | 21.29 (1, 53) | **<0.0001** |
| | **BsnGT** | no treatment | 0.27 ± 0.011 | 16 / 3 | genotype | 91.61 (1, 53) | **<0.0001** |
| | | AP5/CNQX | 0.30 ± 0.013 | 14 / 3 | | | |

**Table 5**

**Relative intensity of phoshoSyn in V1 of WT and BsnEMX1 adult mice 7d after MD**

| Figure | Phosphorylation site | BsnWT 7d MD | | | BsnEmx1 7d MD | | |
|---|---|---|---|---|---|---|---|
| | | Mean ± SEM | | Multiple t test | Mean ± SEM | | Multiple t test |
| | | WT contra | WT ipsi | WT contra vs. ipsi (p-value) | BsnEmx1 contra | BsnEmx1 ipsi | BsnEmx1 contra vs. ipsi (p-value) |
| **Figure 5C** | pSynIa (S9) | 1.312 ± 0.033 | 0.688 ± 0.033 | **<.001** | 0.798 ± 0.009 | 1.202 ± 0.009 | **0,017** |
| | pSynIb (S9) | 1.358 ± 0.034 | 0.642 ± 0.034 | **<.001** | 0.842 ± 0.011 | 1.176 ± 0.011 | 0,124 |
| | pSynIIa (S9) | 1.262 ± 0.130 | 0.738 ± 0.130 | **0,079** | 0.658 ± 0.095 | 1.230 ± 0.026 | 0,055 |
| | pSynIIIa (S9) | 1.265 ± 0.033 | 0.735 ± 0.033 | **0,006** | 0.830 ± 0.064 | 1.170 ± 0.064 | 0,110 |
| | pSynIIb (S9) | 1.186 ± 0.072 | 0.814 ± 0.072 | **0,025** | 0.760 ± 0.031 | 1.240 ± 0.031 | **0,007** |
| | pSynIa (S62) | 1.258 ± 0.055 | 0.742 ± 0.055 | **0,002** | 1.024 ± 0.174 | 1.131 ± 0.038 | 0,417 |
| | pSynIb (S62) | 1.404 ± 0.059 | 0.596 ± 0.059 | **<.001** | 0.986 ± 0.160 | 1.014 ± 0.160 | 0,965 |
| | pSynIIa (S62) | 1.203 ± 0.208 | 0.797 ± 0.208 | 0,197 | 0.793 ± 0.134 | 1.207 ± 0.134 | 0,197 |
| | pSynIIIa (S62) | 1.398 ± 0.146 | 0.602 ± 0.146 | **<.001** | 0.863 ± 0.075 | 1.137 ± 0.075 | 0,204 |
| | pSynIIb (S62) | 1.240 ± 0.118 | 0.803 ± 0.125 | **0,015** | 0.829 ± 0.063 | 1.242 ± 0.061 | **0,020** |
| | pSynIa (S549) | 1.377 ± 0.111 | 0.623 ± 0.111 | **<.001** | 0.878 ± 0.047 | 1.154 ± 0.118 | 0,15 |
| | pSynIb (S549) | 1.382 ± 0.167 | 0.618 ± 0.167 | **<.001** | 0.917 ± 0.051 | 1.083 ± 0.051 | 0,63 |
| | pSynIIa (S549) | 1.483 ± 0.193 | 0.517 ± 0.193 | **<.001** | 0.806 ± 0.147 | 1.194 ± 0.147 | 0,197 |
| | pSynIIIa (S549) | 1.649 ± 0.076 | 0.351 ± 0.076 | **<.001** | 1.120 ± 0.059 | 0.880 ± 0.193 | 0,216 |
| | pSynIIb (S549) | 1.376 ± 0.168 | 0.624 ± 0.168 | **<.001** | 0.997 ± 0.119 | 0.999 ± 0.115 | 0,990 |
| | pSynIa (S553) | 1.305 ± 0.157 | 0.695 ± 0.157 | **<.001** | 0.882 ± 0.137 | 1.118 ± 0.137 | 0,209 |
| | pSynIb (S553) | 1.398 ± 0.120 | 0.602 ± 0.120 | **<.001** | 0.982 ± 0.169 | 1.018 ± 0.169 | 0,506 |
| | pSynIa (S605) | 1.269 ± 0.071 | 0.731 ± 0.071 | **0,001** | 0.890 ± 0.028 | 1.110 ± 0.028 | 0,209 |
| | pSynIb (S605) | 1.431 ± 0.098 | 0.569 ± 0.098 | **<.001** | 0.891 ± 0.052 | 1.109 ± 0.071 | 0,506 |
| | pSynIIa (S605) | 1.171 ± 0.165 | 0.490 ± 0.013 | **0,017** | 0.768 ± 0.168 | 1.073 ± 0.049 | 0,197 |
| | pSynIIIa (S605) | 1.341 ± 0.196 | 0.659 ± 0.196 | **<.001** | 0.972 ± 0.088 | 1.028 ± 0.088 | 0,712 |
| | pSynIIb (S605) | 1.112 ± 0.038 | 0.888 ± 0.038 | 0,193 | 0.799 ± 0.048 | 1.201 ± 0.048 | **0,020** |

**Relative intensity of phoshoSyn in V1 of WT and BsnEMX1 adult mice 7d after sham surgery (no MD)**

| Figure | Phosphorylation site | BsnWT no MD | | | BsnEmx1 no MD | | |
|---|---|---|---|---|---|---|---|
| | | Mean ± SEM | | Multiple t test | Mean ± SEM | | Multiple t test |
| | | WT contra | WT ipsi | WT contra vs. ipsi (p-value) | BsnEmx1 contra | BsnEmx1 ipsi | BsnEmx1 contra vs. ipsi (p-value) |
| **Figure 5C** | pSynIa (S9) | 1.345 ± 0.160 | 0.655 ± 0.160 | 0,096 | 0.986 ± 0.194 | 1.045 ± 0.205 | 0,997 |
| | pSynIb (S9) | 1.318 ± 0.128 | 0.859 ± 0.128 | 0,414 | 1.047 ± 0.199 | 0.953 ± 0.199 | 0,984 |
| | pSynIIa (S9) | 1.341 ± 0.068 | 0.569 ± 0.034 | 0,131 | 0.776 ± 0.166 | 1.205 ± 0.165 | 0,736 |
| | pSynIIIa (S9) | 1.421 ± 0.164 | 0.556 ± 0.142 | **0,023** | 0.808 ± 0.274 | 1.192 ± 0.274 | 0,588 |
| | pSynIIb (S9) | 1.383 ± 0.147 | 0.585 ± 0.116 | **0,002** | 0.908 ± 0.133 | 1.092 ± 0.133 | 0,773 |
| | pSynIa (S62) | 1.223 ± 0.112 | 0.777 ± 0.112 | 0,564 | 0.961 ± 0.183 | 1.039 ± 0.183 | 0,997 |
| | pSynIb (S62) | 1.309 ± 0.170 | 0.858 ± 0.076 | 0,414 | 0.982 ± 0.172 | 1.018 ± 0.172 | 0,984 |
| | pSynIIa (S62) | 1.190 ± 0.258 | 0.810 ± 0.258 | 0,786 | 0.839 ± 0.200 | 1.161 ± 0.200 | 0,800 |
| | pSynIIIa (S62) | 1.235 ± 0.113 | 0.765 ± 0.113 | 0,433 | 0.935 ± 0.171 | 1.065 ± 0.171 | 0,891 |
| | pSynIIb (S62) | 1.359 ± 0.069 | 0.680 ± 0.102 | **0,008** | 1.041 ± 0.109 | 1.237 ± 0.046 | 0,773 |
| | pSynIa (S549) | 1.056 ± 0.054 | 0.944 ± 0.054 | 0,996 | 0.964 ± 0.206 | 1.036 ± 0.206 | 0,997 |
| | pSynIb (S549) | 1.018 ± 0.016 | 0.982 ± 0.016 | 0,984 | 0.876 ± 0.192 | 1.041 ± 0.182 | 0,980 |
| | pSynIIa (S549) | 1.158 ± 0.163 | 0.842 ± 0.163 | 0,800 | 1.171 ± 0.365 | 0.829 ± 0.365 | 0,800 |
| | pSynIIIa (S549) | 1.115 ± 0.142 | 0.885 ± 0.142 | 0,87 | 0.913 ± 0.285 | 1.087 ± 0.285 | 0,891 |
| | pSynIIb (S549) | 1.122 ± 0.167 | 0.878 ± 0.167 | 0,685 | 0.918 ± 0.238 | 1.082 ± 0.238 | 0,773 |
| | pSynIa (S553) | 1.085 ± 0.265 | 0.915 ± 0.265 | 0,993 | 0.962 ± 0.243 | 1.038 ± 0.243 | 0,997 |
| | pSynIb (S553) | 1.171 ± 0.155 | 0.829 ± 0.155 | 0,714 | 0.936 ± 0.253 | 1.064 ± 0.253 | 0,984 |
| | pSynIa (S605) | 1.178 ± 0.099 | 0.822 ± 0.099 | 0,778 | 1.171 ± 0.155 | 1.012 ± 0.196 | 0,993 |
| | pSynIb (S605) | 1.182 ± 0.107 | 0.918 ± 0.013 | 0,873 | 1.041 ± 0.202 | 0.906 ± 0.178 | 0,984 |
| | pSynIIa (S605) | 1.076 ± 0.046 | 0.872 ± 0.078 | 0,800 | 0.933 ± 0.272 | 1.096 ± 0.287 | 0,800 |
| | pSynIIIa (S605) | 1.341 ± 0.196 | 0.659 ± 0.196 | 0,108 | 0.972 ± 0.088 | 1.028 ± 0.088 | 0,891 |
| | pSynIIb (S605) | 1.205 ± 0.057 | 0.795 ± 0.057 | 0,211 | 0.956 ± 0.105 | 0.957 ± 0.042 | 0,997 |

**Table S1**

**Contrast sensitivity (exemplary values at 0.064 cycles / degree)**

| Figure | genotype | treatment / MD-day | Mean ± SEM | Samples | mixed-effects analysis | F(df) | p-value | Mean Contrast |
|---|---|---|---|---|---|---|---|---|
| **Figure S1E** | **BsnWT** | MD / D0 | 15.85 ± 0.56 | 6 | genotype | 2122(1,63) | **<0.0001** | 6,3 |
| P25-35 | | MD / D4 | 22.53 ± 0.86 | 6 | MD-day | 102.5(1,17) | **<0.0001** | 4,4 |
| | **BsnΔEx4-5** | MD / D0 | 2.72 ± 0.06 | 13 | Spatial frequency | 1014(4,68) | **<0.0001** | 36,8 |
| | | MD / D4 | 3.11 ± 0.09 | 13 | Spatial frequency x genotype | 670.6(4,63) | **<0.0001** | 32,2 |
| | | | | | Spatial frequency x MD-day | 37.1(4,63) | **<0.0001** | |
| | | | | | genotype x MD-day | 74.4(1,63) | **<0.0001** | |
| | | | | | Spatial frequency x genotype x MD-da | 33.8(4,63) | **<0.0001** | |
| **Figure S1F** | **BsnWT** | MD / D0 | 15.63 ± 0.97 | 9 | genotype | 67.9(1,44) | **<0.0001** | 6,4 |
| P50-95 | | MD / D7 | 27.97 ± 3.31 | 9 | MD-day | 8.5(1,13) | **0,012** | 3,6 |
| | **BsnΔEx4-5** | MD / D0 | 2.66 ± 0.09 | 6 | Spatial frequency | 84.8(4,52) | **<0.0001** | 37,6 |
| | | MD / D7 | 3.01 ± 0.14 | 6 | Spatial frequency x genotype | 65.0(4,44) | **<0.0001** | 33,2 |
| | | | | | Spatial frequency x MD-day | 9.8(4,44) | **<0.0001** | |
| | | | | | genotype x MD-day | 7.4(1,44) | **0,0094** | |
| | | | | | Spatial frequency x genotype x MD-da | 9.4(4,44) | **<0.0001** | |

**Gain on contrast sensitivity (in %, exemplary values at 0.064 cycles / degree)**

| Figure | genotype | treatment / MD-day | Mean ± SEM | Samples | mixed-effects analysis | F(df) | p-value |
|---|---|---|---|---|---|---|---|
| **Figure S1G** | **BsnWT** | MD / D4 | 43.4 ± 8.0 | 6 | genotype | 14.4(1,17) | **0,0014** |
| P25-35 | **BsnΔEx4-5** | MD / D4 | 14.9 ± 1.8 | 13 | Spatial frequency | 3.1(1.5,23.1) | n.s. |
| | | | | | Spatial frequency x genotype | 2.1(4,63) | n.s. |
| **Figure S1H** | **BsnWT** | MD / D4 | 84.4 ± 22.6 | 9 | genotype | 4.7(1,13) | **0,049** |
| P50-95 | **BsnΔEx4-5** | MD / D4 | 14.0 ± 2.8 | 6 | Spatial frequency | 2.5(1.9,20.4) | n.s. |
| | | | | | Spatial frequency x genotype | 2.7(4,44) | **0,043** |

**Table S2**

**Electrophysiology**

| Figure | genotype | treatment | Mean ± SEM | samples/ independent experiments | statistical test | F(df) | p-value |
|---|---|---|---|---|---|---|---|
| | | | | | **2-way ANOVA** | | |
| **Supplementary figure 2A** | **WT** | no treatment | 1.56 ± 0.09 | 10 / 3 | interaction | 0.1956 (1, 38) | 0,6608 |
| Rise time | | AP5/CNQX | 1.29 ± 0.14 | 12 / 3 | treatment | 2.979 (1, 38) | 0,0925 |
| | **BsnGT** | no treatment | 1.63 ± 0.11 | 10 / 3 | genotype | 1.103 (1, 38) | 0,3002 |
| | | AP5/CNQX | 1.47 ± 0.13 | 10 / 3 | | | |
| **Supplementary figure 2B** | **WT** | no treatment | 8.17 ± 0.30 | 10 / 3 | interaction | 0.3778 (3,76) | 0,7692 |
| Decay time | | AP5/CNQX | 8.58 ± 0.3 | 12 / 3 | treatment | 1.979 (3,76) | 0,1243 |
| | **BsnGT** | no treatment | 7.78 ± 0.52 | 10 / 3 | genotype | 0.04126 (1,76) | 0,8396 |
| | | AP5/CNQX | 8.84 ± 0.56 | 10 / 3 | | | |
| **Supplementary figure 2C** | **WT** | no treatment | 2.91 ± 0.14 | 10 / 3 | interaction | 1.899 (1,38) | 0,1763 |
| Halfwidth | | AP5/CNQX | 2.92 ± 0.14 | 12 / 3 | treatment | 1.930 (1,38) | 0,1729 |
| | **BsnGT** | no treatment | 2.77 ± 0.19 | 10 / 3 | genotype | 0.2323 (1,38) | 0,6326 |
| | | AP5/CNQX | 3.22 ± 0.18 | 10 / 3 | | | |

**Table S3**

**Surface pan-GluA**

| Figure | genotype | treatment | Mean ± SEM | samples/ independent experiments | statistical test | p-value |
|---|---|---|---|---|---|---|
| **Supplementary figure 3A** basal condition | **WT** | no treatment | 1.00 ± 0.07 | 40 / 3 | unpaired t-test | ns |
| | **BsnGT** | no treatment | 0.89 ± 0.05 | 31 / 3 | | |
| **Supplementary figure 3B** | **WT** | no treatment | 1,00 ± 0.07 | 40 / 3 | unpaired t-test | **0,0001** |
| | | AP5/CNQX | 1.52 ± 0.11 | 28 / 3 | | |
| | **BsnGT** | no treatment | 1,00 ± 0.05 | 31 / 3 | unpaired t-test | **<0.0001** |
| | | AP5/CNQX | 1.50 ± 0.10 | 31 / 3 | | |

**Table S4**

| Syt1 uptake assay | genotype | treatment | Mean ± SEM | samples/ independent experiments | 2-way ANOVA | F(df) | p-value |
|---|---|---|---|---|---|---|---|
| **Supplementary fig 4B** | **WT** | no treatment | 1.00 ± 0.06 | 20 / 3 | interaction | 9.936 (1, 85) | **0,0022** |
| Endogenous activity | | AP5/CNQX | 1.42 ± 0.07 | 25 / 3 | treatment | 15.65 (1, 85) | **0,0002** |
| Hippocampal culture | **BsnEmx1** | no treatment | 0.68 ± 0.04 | 21 / 3 | genotype | 73.28 (1, 85) | **<0.0001** |
| | | AP5/CNQX | 0.73± 0.05 | 23 / 3 | | | |
| **Supplementary fig 4D** | **WT** | no treatment | 1.00 ± 0.05 | 26 / 3 | interaction | 4.576 (1, 92) | **0,0351** |
| Evoked activity | | AP5/CNQX | 1.34 ± 0.07 | 27 / 3 | treatment | 15.32 (1, 92) | **0,0002** |
| Hippocampal culture | **BsnEmx1** | no treatment | 0.69 ± 0.04 | 21 / 3 | genotype | 59.18 (1, 92) | **<0.0001** |
| | | AP5/CNQX | 0.79± 0.05 | 22 / 3 | | | |
| **Syt1 uptake assay** | | | | | **2-way ANOVA** | **F(df)** | **p-value** |
| **Supplementary fig 4F** | **WT** | no treatment | 1.00 ± 0.09 | 29 / 3 | interaction | 3.092 (1, 123) | 0,0812 |
| Endogenous activity | | AP5/CNQX | 1.40 ± 0.10 | 34 / 3 | treatment | 12.37 (1, 123) | **0,0006** |
| Cortical culture | **BsnEmx1** | no treatment | 0.63 ± 0.03 | 31 / 3 | genotype | 43.94 (1, 123) | **<0.0001** |
| | | AP5/CNQX | 0.76± 0.04 | 33 / 3 | | | |
| **Supplementary fig 4H** | **WT** | no treatment | 1.00 ± 0.06 | 30 / 3 | interaction | 4.013 (1, 120) | **0,0474** |
| Evoked activity | | AP5/CNQX | 1.35 ± 0.08 | 37 / 3 | treatment | 12.00 (1, 120) | **0,0007** |
| Cortical culture | **BsnEmx1** | no treatment | 0.66 ± 0.05 | 26 / 3 | genotype | 55.22 (1, 120) | **<0.0001** |
| | | AP5/CNQX | 0.75± 0.04 | 31 / 3 | | | |

**Table S4**

**Quantitative western blot analyses of Syn phospho-species in adult mouse V1 following MD**

| Figure | Phosphorylation site | WT 3d MD | | | WT 7d MD | | |
|---|---|---|---|---|---|---|---|
| | | Mean ± SEM | | Multiple t test | Mean ± SEM | | Multiple t test |
| | | WT contra | WT ipsi | WT contra vs. ipsi (p-value) | WT contra | WT ipsi | WT contra vs. ipsi (p-value) |
| **Supl Fig 4** | pSynIa (S9) | 1.205 ± 0.141 | 0.795 ± 0.141 | 0,109 | 1.224 ± 0.017 | 0.776 ± 0.017 | **<.001** |
| | pSynIb (S9) | 1.184 ± 0.110 | 0.816 ± 0.110 | 0,077 | 1.165 ± 0.019 | 0.835 ± 0.019 | **<.001** |
| | pSynIIa (S9) | 1.111 ± 0.053 | 0.889 ± 0.053 | **0,042** | 1.286 ± 0.118 | 0.714 ± 0.118 | **0,027** |
| | pSynIIIa (S9) | 1.162 ± 0.143 | 0.838 ± 0.143 | 0,185 | 1.189 ± 0.093 | 0.811 ± 0.093 | **0,045** |
| | pSynIIb (S9) | 1.253 ± 0.158 | 0.747 ± 0.158 | 0,086 | 1.134 ± 0.038 | 0.866 ± 0.038 | **0,007** |
| | pSynIa (S62) | 1.173 ± 0.071 | 0.827 ± 0.071 | **0,026** | 1.118 ± 0.043 | 0.882 ± 0.043 | **0,018** |
| | pSynIb (S62) | 1.127 ± 0.056 | 0.873 ± 0.056 | **0,033** | 1.128 ± 0.069 | 0.872 ± 0.069 | 0,059 |
| | pSynIIa (S62) | 1.213 ± 0.080 | 0.787 ± 0.080 | **0,02** | 1.181 ± 0.157 | 0.726 ± 0.078 | 0,061 |
| | pSynIIIa (S62) | 1.179 ± 0.076 | 0.821 ± 0.076 | **0,029** | 1.213 ± 0.118 | 0.787 ± 0.118 | 0,062 |
| | pSynIIb (S62) | 1.219 ± 0.088 | 0.781 ± 0.088 | **0,025** | 1.167 ± 0.037 | 0.833 ± 0.037 | **0,003** |
| | pSynIa (S549) | 1.176 ± 0.188 | 0.824 ± 0.188 | 0,257 | 1.198 ± 0.032 | 0.802 ± 0.032 | **<.001** |
| | pSynIb (S549) | 1.172 ± 0.132 | 0.828 ± 0.132 | 0,141 | 1.192 ± 0.065 | 0.808 ± 0.065 | **0,014** |
| | pSynIIa (S549) | 1.268 ± 0.179 | 0.732 ± 0.179 | 0,102 | 1.205 ± 0.069 | 0.795 ± 0.069 | **0,014** |
| | pSynIIIa (S549) | 1.155 ± 0.147 | 0.744 ± 0.140 | 0,113 | 1.171 ± 0.055 | 0.833 ± 0.065 | **0,017** |
| | pSynIIb (S549) | 1.150 ± 0.141 | 0.906 ± 0.146 | 0,296 | 1.177 ± 0.017 | 0.842 ± 0.011 | **<.001** |
| | pSynIa (S553) | 1.123 ± 0.079 | 0.877 ± 0.079 | 0,093 | 1.139 ± 0.070 | 0.810 ± 0.117 | 0,073 |
| | pSynIb (S553) | 1.132 ± 0.090 | 0.868 ± 0.090 | 0,106 | 1.020 ± 0.070 | 0.722 ± 0.197 | 0,227 |
| | pSynIa (S605) | 1.107 ± 0.149 | 0.893 ± 0.149 | 0,370 | 1.135 ± 0.022 | 0.865 ± 0.022 | **0,001** |
| | pSynIb (S605) | 1.104 ± 0.129 | 0.896 ± 0.129 | 0,318 | 1.163 ± 0.068 | 0.837 ± 0.068 | **0,028** |
| | pSynIIa (S605) | 1.056 ± 0.068 | 0.944 ± 0.068 | 0,307 | 1.194 ± 0.077 | 0.806 ± 0.077 | **0,024** |
| | pSynIIIa (S605) | 1.148 ± 0.148 | 0.852 ± 0.148 | 0,232 | 1.217 ± 0.090 | 0.783 ± 0.090 | **0,027** |
| | pSynIIb (S605) | 1.160 ± 0.163 | 0.840 ± 0.163 | 0,237 | 1.173 ± 0.022 | 0.827 ± 0.022 | **<.001** |
